# Molecularly-guided spatial proteomics captures single-cell identity of the healthy and diseased nervous system

**DOI:** 10.1101/2025.02.10.637505

**Authors:** Sayan Dutta, Marion Pang, Gerard M. Coughlin, Sirisha Gudavalli, Michael L. Roukes, Tsui-Fen Chou, Viviana Gradinaru

## Abstract

Single-cell spatial proteomics (scSP) holds substantial potential for profiling healthy and diseased tissues. The emerging method of molecularly-guided unbiased scSP has mostly been applied to peripheral somatic tissues. Here, we optimize and apply scSP to the healthy and diseased mammalian brain, using molecularly-guided laser capture microdissection and unbiased mass spectrometry. We systematically evaluate the effects of tissue fixation, marker staining, and sample input size on proteome coverage and quantitative accuracy. We benchmark this workflow by profiling region-specific neuronal proteomes and describing the response of non-neuronal cells to acute brain injury. Across these applications, we integrate complementary transcriptomic resources to evaluate cross-modality trends and refine neuronal proteomic results by filtering out protein signals likely arising from non-neuronal cells, an essential consideration in heterogeneous tissues such as the brain. Finally, we leverage this approach to resolve proteomic differences between dopaminergic neuron subpopulations with differential vulnerability to Parkinson’s disease and to uncover disease-specific disruptions in α-synuclein-aggregate-bearing single dopaminergic neurons. Together, these data demonstrate the utility of scSP in neuroscience research for understanding fundamental biology and the molecular drivers of neurological conditions.

## Introduction

The central nervous system (CNS) is characterized by molecularly heterogeneous neuronal cells that are spatially organized into distinct subpopulations defined by cell identity and functional states ^1^. Alongside neuronal subpopulations, the mosaic coexistence of glial cells (i.e., astrocytes, microglia, and oligodendrocytes) also contributes to this intrinsic complexity. Resolving such heterogeneity is critical for understanding both the healthy nervous system as well as diseases with sparse pathology^2,3^, requiring approaches that preserve spatial information while enabling molecular profiling at the single-cell resolution. Moreover, human nervous system samples from diseased subjects and small brain regions are often fixed for archiving and available only in limited quantities, making fixation- and single-cell-compatible approaches especially valuable.

Numerous single-cell analysis techniques have been employed to study brain cell types and neurological diseases in a spatially resolved context, although most have focused on transcriptomic insights^4–6^. Protein-focused spatial methods have allowed multiplexed detection of predefined protein targets while preserving spatial context, providing valuable insights into nervous system biology at the single-neuron level^7–9^. Such approaches are particularly powerful for hypothesis-driven analyses, enabling targeted interrogation of known markers across complex tissues and offering functional insights not always evident at the transcript level.

Complementary spatial mass spectrometry (MS)-based methods enable unbiased, discovery-scale proteomic profiling directly from brain tissue and have yielded promising insights in neuroscience^2,10–12^; however, these studies have thus far relied on region-level sampling and have not been applied at single-cell resolution. In parallel, recent advances in MS sensitivity and low-input sample preparation have now made it possible to approach single-cell proteomic measures^13–18^. Recent studies have demonstrated that imaging-guided sample selection, combined with laser-capture microdissection (LCM) or laser ablation and ultrasensitive MS, can recover discovery-scale proteomes with spatial context from individual cells in archival fixed peripheral tissue such as liver and pancreas^19,20^. Despite recent technical advances, molecularly-guided, unbiased single-cell spatial proteomics has not yet been applied to the mammalian nervous system.

Here, we address this gap by presenting a molecularly-guided LCM-based single-cell spatial proteomics (scSP) workflow optimized for paraformaldehyde-fixed brain tissue. We show that unbiased scSP can be applied to interrogate molecular heterogeneity within the mammalian CNS. We explore the potential and constraints of this approach by studying region-specific neuronal identities and describing neuroimmune alterations following focal brain injury. These findings are contextualized within established biological frameworks and through integrative multi-omics analyses, in which single-cell transcriptomic data inform refinement of cellular-origin of proteomic assignments and enable cross-omics comparisons such as regional expression trends between cortex and substantia nigra pars compacta (SNpc). Finally, we apply this platform to define proteomic signatures associated with dopaminergic subtype vulnerability and α-synuclein (αSyn) pathology in a mouse model of Parkinson’s disease (PD), highlighting the broader translational potential of LCM-based scSP for studying disease mechanisms *in vivo*.

## Results

### Molecularly-guided single-cell spatial proteomics (scSP) enables robust proteomic profiling across fixation conditions, staining parameters, and input sample sizes

For unbiased mapping of the proteome from single cells of fixed brain tissue, we adapted^20,21^ a workflow that integrates molecularly-guided selection of micro-tissue samples for downstream proteomic analysis **(Fig. 1A)**. Our workflow begins with tissue fixation, and cryo-sectioning at 30 μm, followed by immunofluorescent (IF) or RNA-fluorescence in-situ hybridization (FISH) staining to label cells of interest, facilitating marker or pathology-guided selection for proteomic analysis. Targeted cells are then precisely isolated using LCM and processed using a sample preparation workflow optimized for low-input, single-cell proteomic analysis^21,22^.

**Figure 1:**
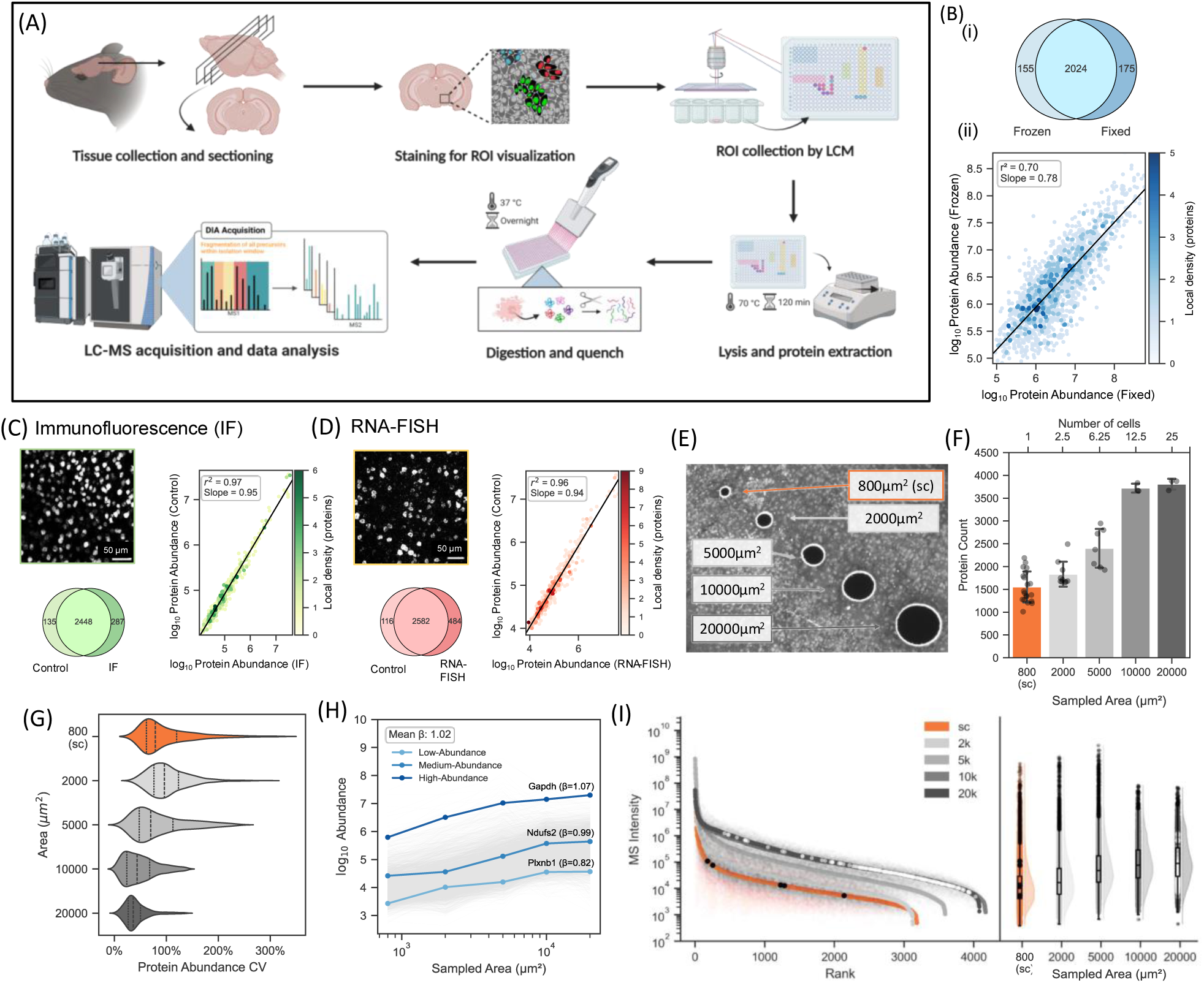
Validation and optimization of marker-guided spatial single-cell proteomics in CNS tissue: **(A)** Schematic overview of the LCM-based scSP workflow. Mouse brain tissue is fixed, cryosectioned, and stained with the desired marker panel, followed by LCM of regions of interest, lysis, digestion, and LC–MS analysis (Created in BioRender. Pang, W. (2025) https://BioRender.com/q97z901). **(B)** Comparison of proteomic profiles obtained from fresh-frozen and fixed bulk cortical tissue, showing protein identification overlap (top) and high correlation of log-transformed protein abundances between conditions (bottom). n = 3 animals **(C)** Effect of IF staining on scSP: representative NeuN-stained mouse cortical section. Scale bar: 50 µm (left, top), overlap of protein identifications between IF-stained samples and unstained (PBS) controls (left, bottom), and correlation of log protein abundances, demonstrating minimal staining-induced bias (right). (n = 10 cells from IF-stained tissue, n = 9 from control tissue) **(D)** Effect of RNA-FISH mRNA staining on protein abundance, showing (i) a representative cortical RNA-FISH image targeting GAD transcripts, (ii) protein identification overlap, and (iii) abundance correlations relative to unstained controls, indicating negligible impact of RNA-FISH on proteomic measurements. (n = 7 cells from FISH-stained tissue, n = 9 cells from control tissue). Scale bar: 50 µm **(E)** Representative NeuN-stained cortical regions corresponding to approximately 1, 3, 6, 12, and 25 neuronal cell bodies (800, 2,000, 5,000, 10,000, and 20,000 µm²; n = 20, 10, 7, 3, and 3, respectively). **(F)** Number of high-confidence proteins (q < 0.01) identified across sampled areas. **(G)** Coefficients of variation (CVs) of protein abundances across sampled areas, showing increased variability at the single-cell level, consistent with greater biological heterogeneity compared to larger cortical regions. **(H)** Log–log scaling of protein abundance as a function of sampled area, with a mean scaling exponent β₁ ≈ 1.02, indicating near-linear scaling across proteins spanning low to high abundance (e.g., Plxnb1, Ndufs2, Gapdh). **(I)** Abundance–rank quantile and raincloud analyses showing proteins uniquely detected in single-cell samples and those absent only from single-cell datasets, with single-cell-specific proteins spanning a broad dynamic range, while proteins missing from single-cell measurements were predominantly low-abundance species.

Before employing stain-guided isolation and processing of samples from fixed tissue, we first assessed whether tissue fixation impacts downstream proteome coverage or introduces bias in detected peptides. Bulk comparison of mouse brain fresh-frozen and fixed tissue, acquired using data-dependent acquisition (DDA), revealed largely overlapping proteome coverage, with 2,024 shared proteins, 175 unique to fixed tissue, and 155 unique to frozen tissue **(Fig. 1B)**. Although statistical testing identified significant differences and the overall log protein abundance correlation was moderate (R^2^ = 0.70; **Fig. 1B, bottom**), no evidence of systematic extraction bias from any cellular compartment was observed between conditions **(Supp. Fig. 1A)**.

We next extended this approach to determine whether IF or RNA-FISH staining protocols introduce measurable proteomic bias. Cortical tissue was stained using anti-NeuN antibody as a representative protein marker **(Fig. 1C)** and FISH probes targeting glutamate decarboxylase 1 and 2 (GAD1+GAD2) mRNA as a representative RNA marker **(Fig. 1D)**.

Comparative analysis was first performed at the bulk proteomic level (using DDA) and subsequently at the single-neuron level using data-independent acquisition (DIA). For both IF-and RNA-FISH-stained samples, bulk proteomes **(Supp. Fig. 1B, 1D)** and single-cell proteomes **(Fig. 1, C, D, Supp. Fig. 1C, 1E)** showed substantial overlap with their respective unstained controls, with comparable protein identifications per cell. Proteins detected uniquely in bulk tissue by IF and RNA-FISH were predominantly low-abundance species, consistent with stochastic sampling effects inherent to DDA rather than systematic staining-related bias **(Supp. Fig. 1B, 1D)**. Log protein abundance values for single-cell analysis were strongly correlated between stained and control samples (R^2^ = 0.97 for IF; R^2^ = 0.96 for HCR), indicating minimal staining-induced bias **(Fig. 1C, 1D)**. Together, these results demonstrate that our IF and RNA-FISH protocols do not introduce systematic bias in proteome coverage or protein abundance and therefore can be confidently employed to define cellular populations within tissues of interest from fixed tissue without compromising the integrity of downstream proteomic measurements.

Next, we evaluated the impact of sample size on proteome coverage and quantification, given that spatial proteomic investigation of the CNS might require samples larger than a single cell in an ROI, or a combination of multiple ROIs. To probe the effect of varying ultra-low input material size, tissue samples corresponding to increasing neuronal cell-body counts were isolated from NeuN stained mice cortex by LCM (due to the finite resolution of cold laser ablation, the effective excised areas were smaller than the nominal cut regions): single cells (∼800 µm^2^), ∼3 cells (2,000 µm^2^), ∼6 cells (5,000 µm^2^), ∼12 cells (10,000 µm^2^), and ∼25 cells (20,000 µm^2^) **(Fig. 1E, Supp. Fig. 2A-C)**. All samples were analyzed using a previously optimized 1-hour LC-MS gradient with DIA^22,23^, which was specifically tuned for single-cell inputs. Notably, single-cell samples (n = 20 single cells) yielded consistently high proteome depth, with up to ∼2,200 proteins quantified from an individual neuron and protein identifications increased with sampled area but plateaued beyond 10,000 µm^2^ **(Fig. 1F)**. A similar pattern was observed for the peptide quantification **(Supp. Fig. 2D)**. Higher coefficients of variation (CVs) in single-cell samples relative to larger sampled areas from the cortex **(Fig. 1G)** likely reflects biological heterogeneity unmasked at single-cell resolution.

**Figure 2:**
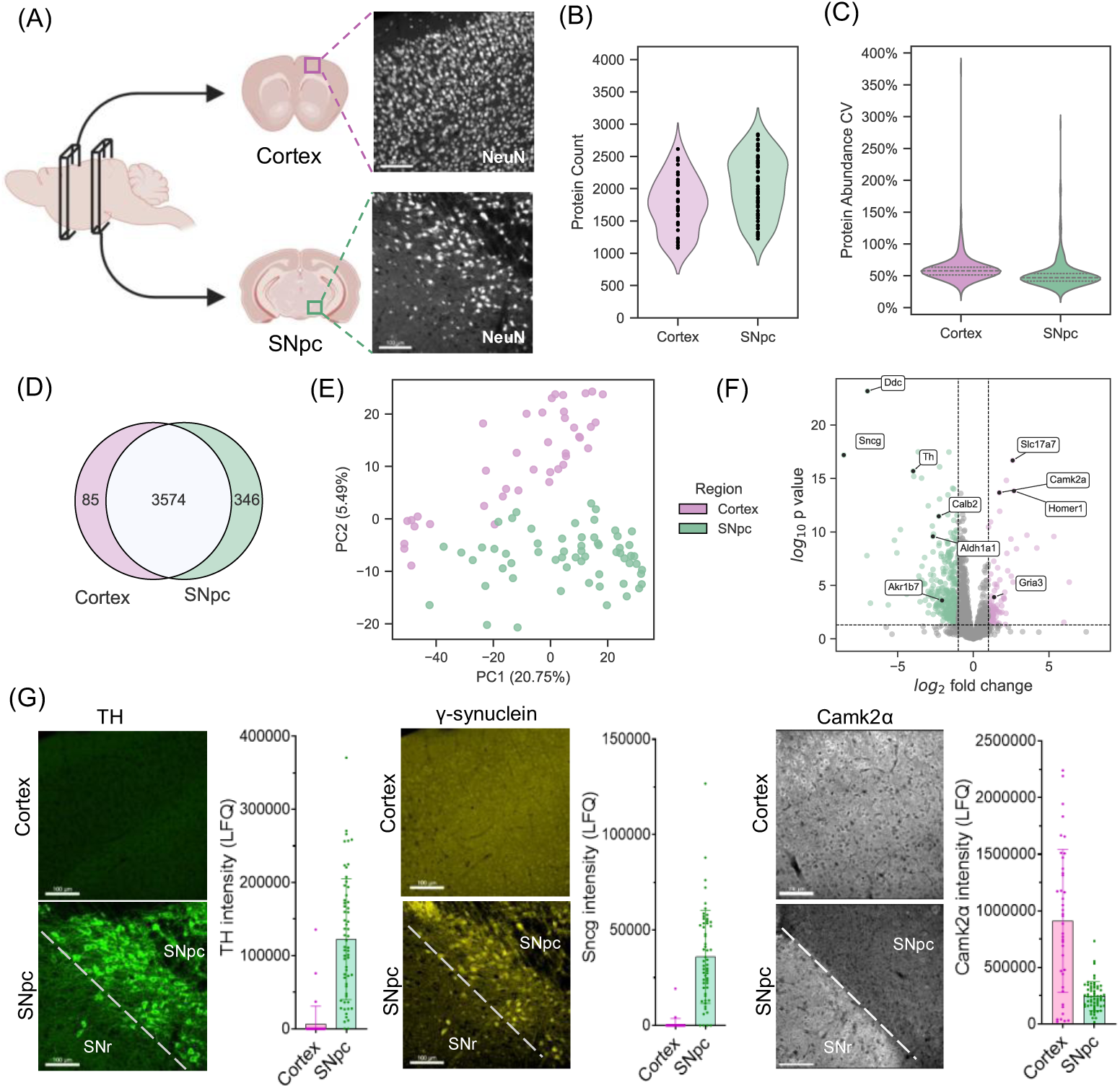
Characterization of CNS neuronal subpopulations using LCM-based scSP: **(A)** Schematic representation of the cortex and SNpc brain regions selected for comparison using LCM-based scSP. Representative confocal microscopy images showing immunostaining of neuronal cell bodies in the cortex and SNpc. Neurons are labeled with anti-NeuN. Created in BioRender. Dutta, S. (2026) https://BioRender.com/t68m928 (modified). Scale bar: 100 µm. **(B)** Protein counts from isolated single-cell samples obtained from cortical and SNpc regions using LCM-based scSP. (n= 60 cells for cortex; 40 cells for SNpc). **(C)** CVs of protein abundances for the groups showing comparable variance of around 50% for both groups. **(D)** Overlap of identified proteins between cortical and SNpc samples and proteins uniquely identified in each group. **(E)** Unsupervised PCA plot of protein intensities from the two sample groups showing cluster separation of single-cell samples from the cortex and SNpc. **(F)** Distribution of differentially expressed proteins (DEPs) between the cortex and SNpc. Proteins significantly upregulated in each neuron type are highlighted, revealing region-specific protein markers. **(G)** Corresponding IF validation using anti-TH, anti-gamma synuclein, and anti-Camk2α antibodies to indicate protein expression distribution. The respective bar graphs illustrate the label-free quantification (LFQ) mass spectrometry intensity levels of these marker proteins in each single-cell sample.

Because single-cell measurements operate in a lower signal-to-noise regime than multi-cell samples, we next assessed whether protein abundance scales proportionally with sampled area. Protein abundance was modeled using a log-log scaling relationship, where the slope 𝛽_1_ represents the scaling exponent. After removal of contaminants **(Supp. Fig. 2E)**, the mean value of 𝛽_1_ across all proteins (gray lines) was 1.02 **(Fig. 1H)**, indicating near-linear scaling between sampled area and measured abundance. This behavior was consistent across low-, medium-, and high-abundance proteins (e.g., Plxnb1, Ndufs2, and Gapdh, respectively; **Fig. 1H)**. In contrast, contaminants such as keratins exhibited high variability across sample sizes **(Supp. Fig. 2, E-F)**, consistent with sample preparation artifacts rather than biological effects. Together, these results indicate that, after accounting for contaminants, protein abundance measurements scale proportionally with input material and are not detectably biased by sample size.

Finally, we examined whether single-cell sampling selectively biases protein detection. Analysis of proteins across different sample areas revealed that only 5 proteins were uniquely detected in single-cell samples, whereas 182 proteins were consistently detected in all larger sampled areas but absent from single-cell measurements **(Supp. Fig. S2G)**. No significant functional enrichment was observed among the proteins specific to single cells. An abundance-rank plot shows the distribution of these proteins **(Fig. 1I)**, with single-cell-specific proteins spanning a wide dynamic range, indicating that single-cell samples can detect a broad range of proteins without abundance bias. Proteins missing from single-cell samples (shown in white) were predominantly low-abundance proteins, consistent with the expectation that these proteins fall below detection thresholds in single-cell samples due to MS sensitivity limitations. Therefore, although proteome coverage at the single-cell level remains limited by low signal-to-noise, the use of DIA mitigates stochastic sampling effects relative to DDA. Collectively, these findings establish LCM-based scSP as a robust and quantitatively unbiased approach for proteomic characterization of individual neurons, motivating its application to biologically and spatially defined neuronal populations in subsequent analyses.

### Single-cell spatial proteomics reveals region-specific neuronal proteomes

We applied LCM-based scSP to characterize neuronal subtypes from distinct brain regions **(Fig. 2A)**. PFA-fixed coronal brain slices from adult mice were immunostained with the pan-neuronal marker anti-NeuN to enable molecularly-guided micro-dissection of single neuronal cell bodies from the cortex (approximately from layers IV-V of the primary motor cortex) and the substantia nigra pars compacta (SNpc) for subsequent proteomic profiling. We identified approximately 2,000 proteins on average per cell (n= 40 cortex, 60 SNpc single cells; **Fig. 2B**). Violin plots show comparable protein abundance CV distributions between cortex and SNpc, with median CV values centered around ∼50–60% in cortex and ∼45–50% in SNpc, and extended distribution tails reflecting the presence of highly variable proteins **(Fig. 2C)**. We identified 3574 protein groups present in both neuronal populations, with an additional 85 proteins exclusively detected in cortical neurons and 346 in SNpc neurons **(Fig. 2D)**. Proteins detected exclusively in one region may reflect biologically meaningful regional specificity. For example, Icam5 and Slc6a3 (DAT), proteins known to be highly expressed in cortical neurons and SNpc, respectively, were identified in the cortex-specific and SNpc-specific protein sets.

Unsupervised principal component analysis (PCA) revealed two distinct clusters, highlighting significant molecular differences between cortical and SNpc neurons **(Fig. 2E)**. Among the 3574 proteins identified in both groups, 99 proteins exhibited more than two-fold upregulation in cortical neurons and 345 proteins in SNpc neurons. This analysis identified multiple regionally selective proteins, including Camk2a, Gria3, Homer1, and Slc17a7 in cortical neurons, whereas SNpc neurons were enriched for Th, Ddc, Sncg, Calb2, Aldh1a1, and Akr1b7. To validate these proteomic findings, we cross-referenced some of the identified proteins with in-situ hybridization (ISH) gene expression data from the Allen Brain Atlas^24^ **(Supp. Fig. 3)** and performed IF validation **(Fig. 2G)**. Additionally, we plotted MS-derived intensity values, which showcase the heterogeneity in protein expression at the individual cell level **(Fig. 2G)**. Together, these results demonstrate that LCM-based scSP robustly resolves region-specific neuronal proteomes at single-cell resolution.

**Figure 3:**
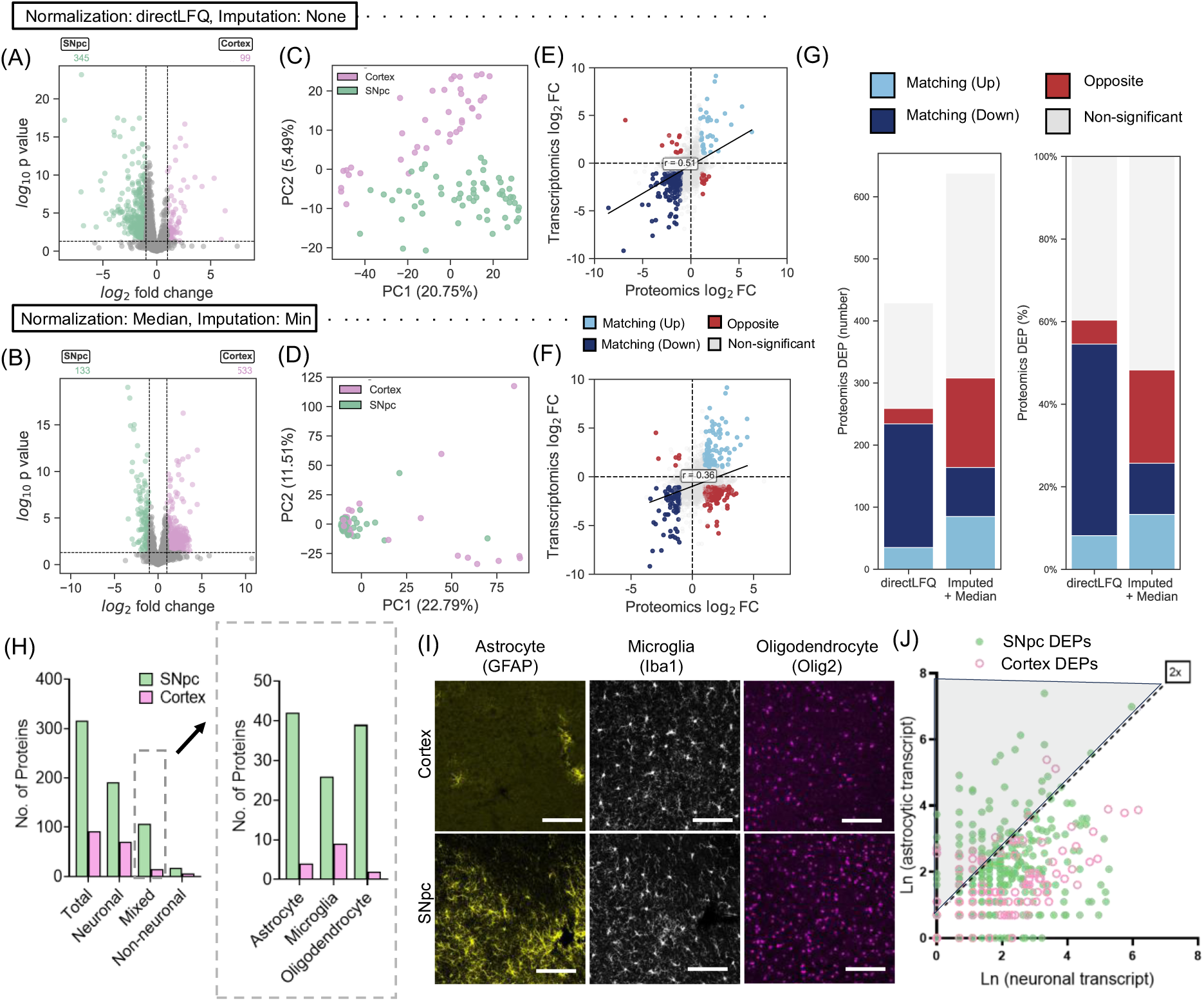
Single-cell transcriptomics informs data processing and refinement of neuronal proteomic assignments. (A–B) Distribution of DEPs between cortical and SNpc neurons using directLFQ quantification (A) or imputation (20% minimum value) followed by global median normalization (B), highlighting increased DEP detection and cortex-skewed upregulation with imputation. **(C–D)** Unsupervised PCA of single-neuron proteomes demonstrates improved separation of cortical and SNpc samples using directLFQ (C) compared to imputed data (D). **(E–F)** Cross-omics comparison of proteomic log₂ fold changes with single-cell transcriptomic cortex-to-SNpc ratios, color-coded by concordant upregulation, concordant downregulation, and opposite directionality, reveals higher concordance and correlation using directLFQ. **(G)** Quantification of agreement between proteomic and transcriptomic directionality shows higher overall concordance for directLFQ relative to imputed normalization. **(H)** Classification of DEPs based on Drop-seq transcriptomic enrichment using DropViz (i) classifies proteins as neuronal, mixed, or non-neuronal. (ii) Stratification of mixed DEPs into confidence groups based on the highest transcript levels among astrocytes, microglia, and oligodendrocytes, possible contributors to background signal in SNpc proteomes. **(I)** IF staining of astrocytes, microglia, and oligodendrocytes reveals greater astrocytic density in SNpc compared to cortex, confirming astrocytes as major contributors to non-neuronal DEPs in SNpc samples. Scale bar 100 μm. (J) Correlation plot of neuronal and astrocytic transcript abundance for DEPs visualizes cell-type-specific expression patterns. Gray shaded area indicates DEPs of possible astrocytic origin.

### Single-cell transcriptomics informs data processing and refinement of neuronal proteomic assignments

Single-cell-level proteomics lacks standardized processing and normalization workflows^25,26^. In addition to the presence of substantial missing values within datasets due to the minute amount of starting material, single-cell measurements are characterized by reduced signal-to-noise ratios, making quantitative comparisons particularly sensitive to pre-processing strategies. To address this issue, various normalization and data imputation strategies are often applied during analysis^27–29^. Here, we compared peptide-level directLFQ normalization^30^ with a protein-level median-based normalization strategy incorporating minimum-value imputation (20% of minimum) to evaluate whether one approach performs better on single-cell spatial proteomics datasets. The median-based normalization approach incorporating imputation increased the number of differentially expressed proteins (DEPs) identified between cortex and SNpc relative to directLFQ and produced a skewed distribution, with more proteins classified as upregulated in cortex **(Fig. 3, A-B)**. However, unsupervised PCA demonstrated improved separation of cortical and SNpc samples using directLFQ compared to the median-based normalization approach **(Fig. 3, C-D)**. Although many established regional markers showed concordant regulation across both approaches, a subset of proteins exhibited discordant directionality. To resolve this conflict, we used cross-omics comparison with single-cell transcriptomics datasets to assess these two normalization strategies. For this, we used the 10x scRNA-seq dataset of the mouse whole-brain transcriptomic dataset from Allen Brain Atlas^31^ and extracted the dopaminergic neurons within the midbrain transcriptomic dataset, corresponding to our SNpc proteomics cells, and IT–ET (intratelencephalic and extratelencephalic) glutamatergic neurons within the MOp (primary motor cortex) dataset, corresponding to our cortical proteomics cells. We observed clear separation of the cell clusters and identification of region-specific enriched genes **(Supp. Fig. 4)**, some of which we also identified and validated in our proteomics datasets **(Fig. 2)**.

**Figure 4.**
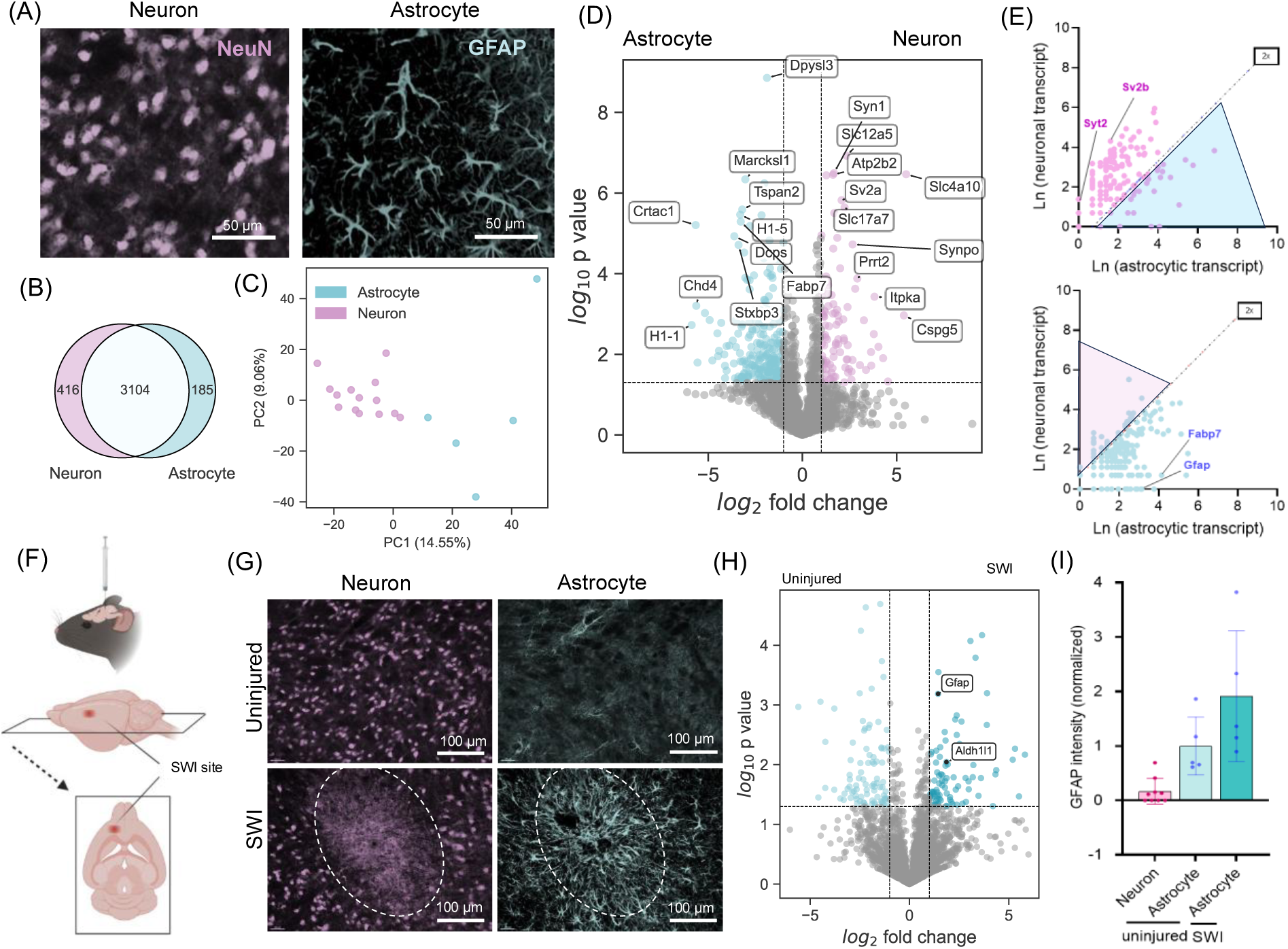
Single-cell spatial proteomics of astrocytes capture cell-type identity and neuroinflammatory activation. **(A)** Representative IF images of mouse brain sections stained for NeuN (neurons) and GFAP (astrocytes), enabling selective laser capture. Scale bars, 50 µm. **(B)** Venn diagram showing overlap and cell-type-specific protein groups identified across single neurons and astrocytes. (n= 14 cells for neuron; 5 cells for astrocyte) **(C)** Unsupervised PCA of single-cell proteomic profiles demonstrates clear separation between neuronal and astrocytic samples. **(D)** Volcano plot depicting differentially abundant proteins between astrocytes and neurons. Representative neuron-enriched and astrocyte-enriched proteins are annotated. **(E)** Integration of proteomic differential abundance with Drop-seq transcriptomic data, comparing astrocyte- and neuron-enriched proteins against corresponding astrocytic and neuronal transcript expression, respectively (shaded area highlights potential non-specific DEPs). **(F)** Schematic of the acute stab wound injury (SWI) model, illustrating the injury site. Created in BioRender. Dutta, S. (2026) https://BioRender.com/kye7fc7 (modified). **(G)** Representative IF images of NeuN and GFAP in uninjured control tissue and at the SWI site. Scale bars, 100 µm. **(H)** Differential abundance analysis of single astrocytes isolated from the peri-lesional SWI region compared with astrocytes from contralateral uninjured tissue (n= 5 cells for uninjured; 5 cells for SWI). Proteins associated with astrocyte activation are highlighted. **(I)** Quantification of mass spectrometry-derived GFAP intensities across neurons and astrocytes from uninjured and SWI conditions, showing increased GFAP abundance in reactive astrocytes.

We compared cortex-to-SNpc transcript ratios derived from single-cell transcriptomic data with proteomic log2 fold changes, focusing on directionality rather than effect size, as a pragmatic benchmark for evaluating normalization strategies **(Supp. Table 2)**. While transcript-protein correlations are known to be limited at single-cell resolution due to post-transcriptional regulation, protein stability, temporal delays, and stochastic gene expression^32–34^, consistent regional trends across modalities provide a useful reference for assessing whether proteomic processing preserves biologically plausible signal structure. Using directLFQ normalization, we identified 234 gene-protein pairs with consistent up- or down-regulation and 25 pairs with opposite directionality, corresponding to ∼80% directional agreement and a higher correlation (R² = 0.51). In contrast, the median-based normalization approach incorporating minimum-value imputation yielded 164 same-direction and 144 opposite-direction pairs, corresponding to ∼40% agreement and a lower correlation (R² = 0.36) **(Fig. 3, E-G)**. These results highlight that normalization strategy substantially influences downstream biological interpretation, and that leveraging orthogonal omics data provides a principled framework for evaluating single-cell proteomics processing methods. Consequently, we adopted directLFQ as the normalization approach for all subsequent analyses.

Closer inspection of a small subset of proteins (∼5.8%) that show disagreement in single-cell transcriptomic directionality following directLFQ normalization **(Fig. 3G)**, revealed that several corresponded to proteins of non-neuronal origin, including MBP (oligodendrocyte marker) and GFAP (astrocytic marker). Because our isolation strategy relied on laser microdissection rather than membrane-enclosed methods for intact single cells (i.e., cultured cells or separation by cell sorting), complete exclusion of proteome contributions from neighboring cells is impossible to achieve. These contaminating proteins can confound investigation of cell type-specific proteomes. To systematically address this issue, we integrated single-cell transcriptomic data from a publicly available Drop-seq mouse dataset (Dropviz)^35^ to assign a probable cellular origin to the DEPs (**Supp. Table 3**). Proteins with higher neuronal transcript counts, including those exclusively detected in neurons, or with neuronal counts within 50% of any non-neuronal cell type (empirically defined threshold, given the LCM isolation is neuron-targated) were classified as ‘neuronal’. Proteins with zero neuronal transcript counts were flagged as exclusively ‘non-neuronal’ (17 in SNpc and 6 in cortex; **Fig. 3H, left**). DEPs with transcripts detected in both neuronal and non-neuronal cells, but with neuronal counts less than 50% of any of the non-neuronal cell types, were classified as mixed and assigned lower confidence due to ambiguity in assigning cell-type origin. Interestingly, ∼39% of SNpc DEPs were classified as ‘mixed’ or ’non-neuronal’, compared to ∼22% for cortical DEPs, suggesting a greater infiltration of one or more non-neuronal cell types within the SNpc neuronal regions compared to the cortex. In ’mixed’ DEPs, we observed substantial mRNA counts across multiple non-neuronal cell types, including astrocytes, microglial and oligodendrocytes, preventing confident assignment to a single non-neuronal class **(Fig. 3H, right)**. However, parallel IF results **(Fig. 3I)** identified that astrocytes were the most disproportionately enriched non-neuronal population in SNpc relative to cortex, implicating them as the primary contributor to the non-neuronal component of the SNpc proteome DEP hits. Plotting the correlation of transcriptomic abundance of DEPs in neurons and astrocytes further supports this (**Fig. 3J**, gray area indicates DEPs of ‘mixed’ and ‘non-neuronal’ origin).

Taken together, these results indicate that single-cell transcriptomics can be integrated with scSP both to evaluate differential expression trends across modalities and to probabilistically assign cellular origin to proteomic signals, an especially critical consideration in complex tissues such as the CNS. In subsequent analyses, this framework is leveraged to refine cell-type attribution and biological interpretation of proteomic changes across multiple spatial and cellular analyses.

### Single-cell spatial proteomics of non-neuronal cells in the CNS can capture markers of neuroinflammation

Glial cells are essential for maintaining CNS homeostasis, providing structural support, regulating neuronal function, and modulating inflammatory responses^36,37^. Given their importance, we next asked whether non-neuronal cell types could be reliably distinguished and profiled using our approach, focusing initially on astrocytes. Brain sections were immunostained with the astrocytic marker GFAP together with NeuN to enable selective isolation of non-overlapping astrocyte and neuron populations **(Fig. 4A)**. Across both cell types, we identified 3,104 protein groups, including 185 proteins uniquely detected in astrocytes and 416 uniquely detected in neurons **(Fig. 4B)**. PCA revealed cell-type-specific clustering of astrocytes and neurons **(Fig. 4C)**. Differential abundance analysis identified multiple lineage-specific proteins **(Fig. 4D)**: neuron-enriched proteins included Bsn, Sncb, Syn1, and Dlg4, whereas astrocyte-enriched proteins included Fabp7, Vim, Anxa2, and GFAP. Integration with transcriptomic datasets confirmed that neuronal DEPs corresponded predominantly to neuron-enriched transcripts, while astrocyte-upregulated DEPs showed enrichment in astrocytic transcripts **(Fig. 4E)**.

We next applied LCM-based scSP to investigate neuroinflammation following acute brain injury using a stab wound injury (SWI) model^38^, which induces rapid neuronal loss along the needle track and formation of a surrounding glial scar **(Fig. 4, F-G)**. GFAP⁺ astrocytes were isolated from the peri-lesional region and compared with GFAP⁺ astrocytes from the contralateral, uninjured hemisphere. Both groups had comparable protein counts per cell, with 422 proteins uniquely detected in the injured side and 512 in the uninjured side. Unsupervised PCA revealed separation between injured and control astrocytes **(Supp. Fig 5, A-C)**. Proteomic analysis revealed significant upregulation of proteins associated with astrocytic activation, including Aldh1l1 and GFAP **(Fig. 4H)**, with increased GFAP abundance evident at the single-sample level **(Fig. 4I)**.

**Figure 5:**
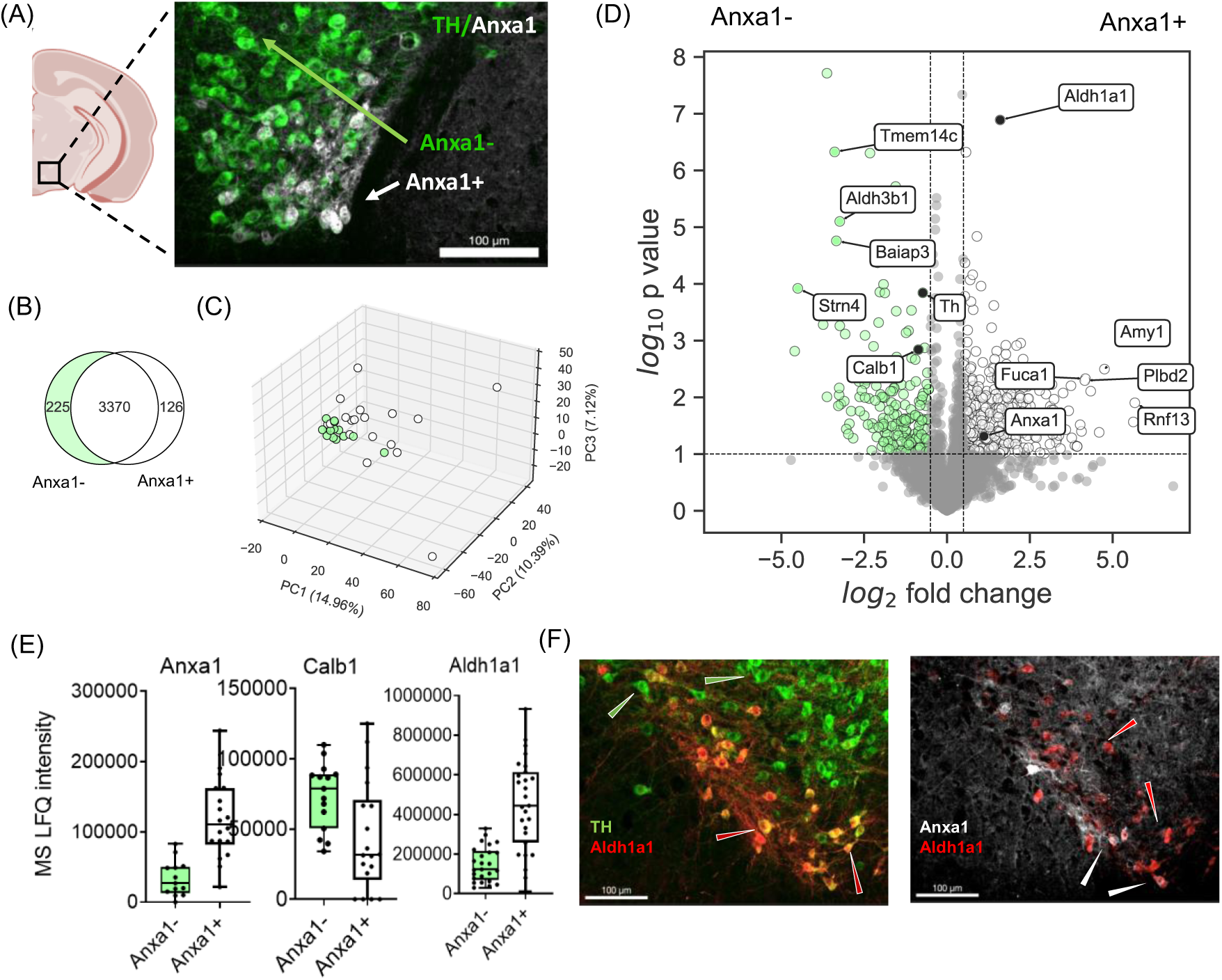
Proteomic differences between dorsal and ventral SNpc dopaminergic neuron subpopulations determined by scSP. **(A)** Schematic and representative IF image illustrating dual labeling of SNpc dopaminergic neurons with TH (green) and Anxa1 (white) to distinguish ventral-tier (TH⁺/Anxa1⁺) from dorsal-tier (TH⁺/Anxa1⁻) neurons. Scale bar, 100 µm. Created in BioRender. Dutta, S. (2026) https://BioRender.com/t5i05fp (modified). **(B)** Venn diagram showing overlap of identified protein groups across TH⁺/Anxa1⁺ and TH⁺/Anxa1⁻ single neurons, with population-specific proteins indicated (n = 18 cells for Anxa+; 15 cells for Anxa-). **(C)** PCA of profiles from Anxa1⁺ and Anxa1⁻ dopaminergic neurons **(D)** Differential protein abundance between Anxa1⁺ and Anxa1⁻ neurons, highlighting top proteins enriched in each sample group. **(E)** Protein abundance distributions for Anxa1, Calb1, and Aldh1a1 in Anxa1⁺ and Anxa1⁻ neurons, illustrating differential expression between the groups. **(F)** Representative IHC validation of higher expression of Aldh1a1 in the ventral SNpc neurons and lower or absent staining in the dorsal tier. Scale bar, 100 µm.

Notably, acute SWI induces robust microglial activation and ramification near the injury site **(Supp. Fig. 5D)**, and we detected several microglial inflammatory markers (e.g., AIF1, CD68, CD11b, C1qa) in the micro-dissected astrocytic samples, which we validated via IF **(Supp. Fig. 5, E-F)**, indicating that microglial populations could be amenable to LCM-based scSP. We therefore attempted to profile microglia in control brain tissue using AIF1/Iba1 labeling. However, this proved technically challenging, likely due to the small size of microglial cell bodies, necessitating methodological refinements such as thinner tissue sections and reduced LCM capture areas.

Together, these results demonstrate that LCM-based scSP enables robust, cell-type-resolved proteomic profiling of glial populations *in situ* and provides a powerful framework for studying region-specific neuroinflammatory responses, including reactive gliosis, an essential pathological feature of stroke, traumatic brain injury, and neurodegenerative disease.

### Proteomic profiling of Parkinson’s disease-vulnerable ventral SNpc neurons reveals basal molecular differences within closely related dopaminergic subpopulations

Building on our cortex-SNpc comparison, we next asked whether LCM-based scSP could resolve proteomic differences between molecularly defined neuronal populations that are highly similar in anatomy and function and inaccessible to bulk approaches. We focused on dorsal and ventral neuronal subpopulations of SNpc, the latter of which selectively degenerate in PD^4,39^. To isolate these populations, we employed dual immunostaining for TH to label SNpc dopaminergic neurons and annexin A1 (Anxa1) to selectively mark the ventral tier population^40^, enabling comparison of TH⁺/Anxa1⁺ ventral neurons with TH⁺/Anxa1⁻ dorsal neurons **(Fig. 5A)**. We identified 3,370 protein groups common to both Anxa1⁺ and Anxa1⁻ neurons, with an additional 126 proteins uniquely detected in Anxa1⁺ neurons and 225 uniquely detected in Anxa1⁻ neurons (n = 15–17 single neurons per group; **Fig. 5B**). PCA failed to robustly separate the two populations, possibly due to their anatomical and functional proximity **(Fig. 5C)**.

Therefore, we used relaxed statistical thresholds (p < 0.1, FC ≥ 1.5) to identify 370 proteins significantly upregulated in Anxa1⁺ neurons and 189 enriched in Anxa1⁻ neurons **(Fig. 5D)**. The dataset was enriched for canonical neuronal proteins, including synaptic, cytoskeletal, and neurotransmission-related markers such as Map2, Camk2a, Th, Stmn3, Vamp1, Cadps2, Calb1, and Aldh1a1, confirming neuronal identity at the proteomic level. Within the neuronal proteome, Anxa1 itself was differentially regulated, as expected **(Fig. 5E)**. Notably, Aldh1a1 was enriched in ventral-tier neurons, whereas Calb1 was elevated in dorsal-tier neurons; these proteins have been previously implicated in dopamine metabolism and calcium homeostasis, respectively, and could be associated with differential vulnerability of SNpc subpopulations **(Fig. 5E)**. These findings were validated by IF **(Fig. 5F, Supp. Fig. 6)** and are consistent with prior trascriptomics reports^40,41^. Together, these results demonstrate that LCM-based scSP can resolve subtle yet biologically meaningful proteomic differences between closely related neuronal subpopulations, providing complementary insight to transcriptomic approaches^42^ for dissecting cell type-specific mechanisms underlying selective neuronal vulnerability in PD and related disorders.

**Figure 6.**
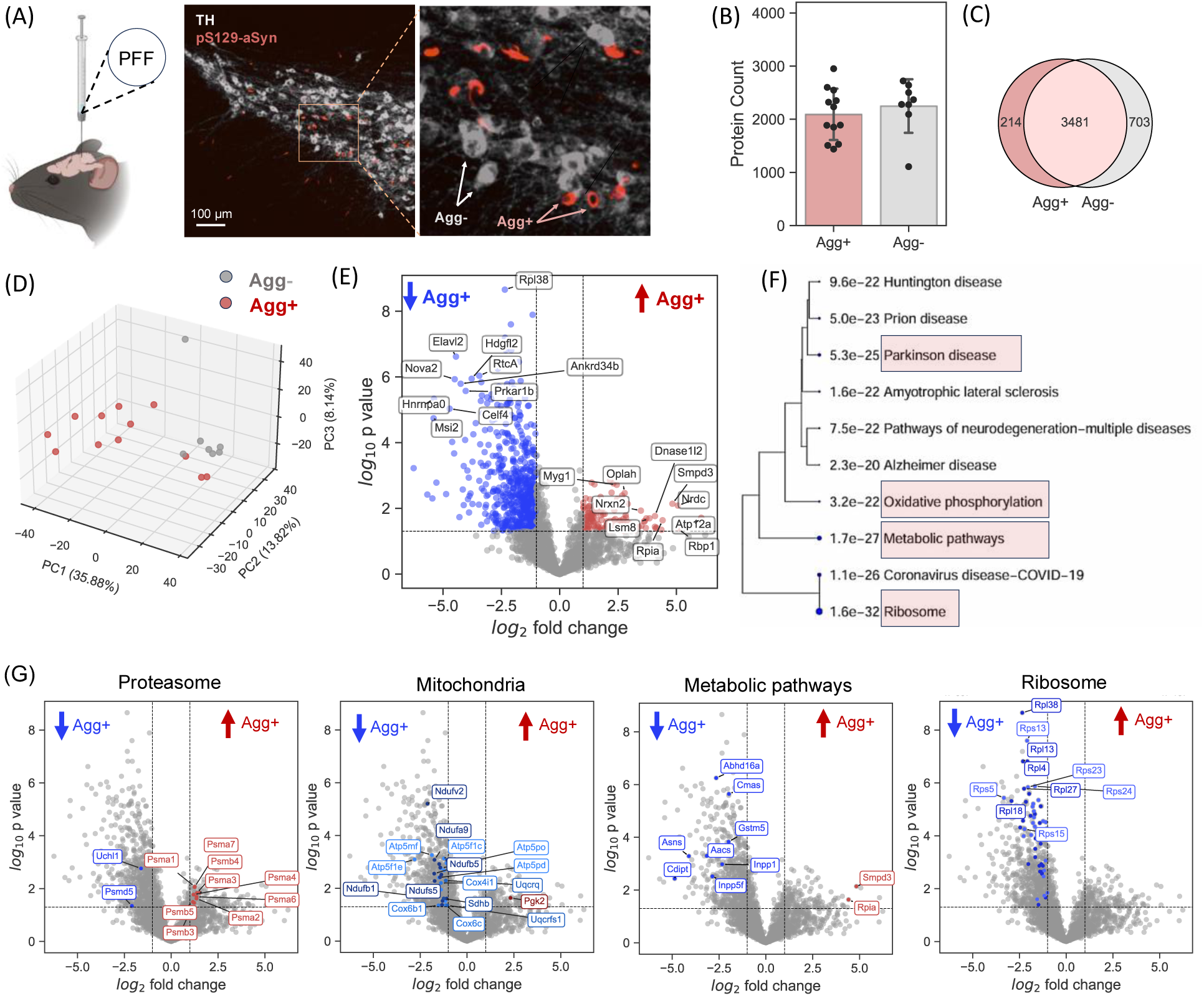
Proteomic profiling of α-synuclein aggregate–bearing SNpc neurons in a murine Parkinson’s disease model. **(A)** Experimental overview and representative IF showing dopaminergic SNpc neurons labeled with TH and pathogenic phosphorylated α-synuclein (pS129-αSyn) aggregates, enabling identification and laser capture of aggregate-containing (Agg⁺) and non-aggregate (Agg⁻) neurons following intrastriatal injection of αSyn preformed fibrils (PFFs). Created in BioRender. Dutta, S. (2026) https://BioRender.com/s7wcjyf (modified). **(B)** Number of proteins identified per single neuron in Agg⁺ and Agg⁻ populations (mean ± SD). (n= 12 cells for Agg+; 7 cells for Agg-) **(C)** Venn diagram showing overlapping and uniquely detected proteins between Agg⁺ and Agg⁻ neurons. **(D)** Unsupervised PCA of single-cell proteomes, illustrating separation and increased variability of Agg⁺ neurons. **(E)** Volcano plot of differential protein abundance between Agg⁺ and Agg⁻ neurons (log₂ fold change versus –log₁₀ p value), with selected differentially abundant proteins annotated. **(F)** KEGG pathway enrichment analysis of differentially abundant proteins showing top 10 implicated pathways. **(G)** Pathway-focused volcano plots illustrating differential regulation of proteins associated with the proteasome, mitochondrial oxidative phosphorylation, metabolic pathways, and the ribosome in Agg⁺ neurons relative to Agg⁻ neurons.

### Single-cell spatial proteomics identifies proteins and pathways altered in pS129-αSyn inclusion-bearing neurons in a murine model of Parkinson’s disease

Deciphering molecular-level insights selectively from the small, spatially restricted neuronal populations that harbor pathogenic protein inclusions in neurodegenerative disorders such as PD remains challenging. For example, in human post-mortem samples from patients with PD or dementia with Lewy bodies, only ∼1-2% of neurons contain mature Lewy bodies^3^, and even in inducible murine models, inclusion-bearing neurons typically comprise only a fraction of the local SNpc population^43^. A recent study used spatial transcriptomics to map molecular alterations in pathology-enriched brain regions^44^; however, proteomic changes at the single-cell level have not been investigated. To address this, we applied LCM-based scSP to a murine model of PD, modeling the symptomatic stage of parkinsonism using direct injection of pathogenic αSyn preformed fibrils (PFFs)^45–47^. Adult C57 mice received intrastriatal injections of sonicated mouse PFFs, and brain tissue was collected 6 months later. IHC confirmed αSyn aggregates marked by pS129-αSyn in a fraction of surviving SNpc neurons **(Fig. 6A)**, and these inclusions also stained positive for canonical pathology markers such as ubiquitin and p62 **(Supp. Fig. 7)**.

Aggregate-containing (Agg⁺) and non-aggregate-containing (Agg⁻) neurons **(Fig. 6A)** were laser-captured from PFF-injected mice nigral tissue dual-labeled for a dopaminergic marker (TH) and anti-pS129-αSyn antibody, with non-aggregate neurons collected from the PBS-injected contralateral control side of the same animals. On average, we detected ∼2,000 unique proteins per individual neuron **(Fig. 6B)** and ∼4300 protein groups across both neuronal populations, with 703 proteins exclusive to Agg⁻ neurons and 214 proteins to Agg⁺ neurons **(Fig. 6C)**. Unsupervised dimensionality reduction resolved the two neuronal groups into distinct clusters, with Agg⁺ neurons showing markedly higher variability **(Fig. 6D)**. Analysis of DEPs **(Fig. 6E)** and pathway enrichment analysis of the DEPs using KEGG, a curated compendium of experimentally validated molecular networks^48^, revealed strong overrepresentation of pathways related to “Parkinson’s disease,” “Oxidative phosphorylation,” “Metabolic pathways,” and the “Ribosome” **(Fig. 6F)**. Consistently, gene ontology enrichment across cellular components, molecular functions, and biological processes yielded overlapping categories **(Supp. Fig. 8)**.

Mapping the altered proteins onto the KEGG Parkinson’s disease pathway highlighted multiple nodes along established pathogenic routes **(Supp. Fig. 9)**. Dysregulation of the ubiquitin-proteasome system was evident, marked by increased abundance of several proteasomal subunits (e.g., PSMB5, PSMA7) in Agg⁺ neurons together with reduced levels of the familial PD-linked deubiquitinase UCHL1 **(Fig. 6G)**. Mitochondrial dysfunction also emerged prominently, reflected by widespread downregulation of subunits from respiratory chain complexes I-V. Notably, numerous ribosomal proteins (RPL and RPS families) were decreased in Agg⁺ neurons. Network analysis further identified a connected module of ribosome-associated RNA-binding proteins, including several spliceosome components **(Supp. Fig. 10)**, suggesting coordinated perturbation of translational and RNA-processing machinery. These results support multi-pathway alterations in αSyn aggregate-containing neurons in this model of Parkinsonism.

Together, these findings illustrate how LCM-based scSP enables proteome-wide interrogation of rare, pathology-defined neuronal subpopulations *in vivo*, offering a powerful framework to single-cell resolution analysis of proteomic states associated with neurodegenerative diseases.

## Discussion

Laser capture microdissection-based spatial proteomics has emerged as a key method for studying complex biological systems, enabling interrogation of deep proteomic complexity while preserving tissue context^49,50^. To date, spatial proteomics at the single-cell level has only been applied to peripheral somatic tissues with comparatively lower cellular diversity or spatial complexity^19,51^, and application to the brain remains challenging due to its cellular and architectural complexity. Here, we apply LCM-based single-cell spatial proteomics (scSP) to multiple neuroscience use cases, demonstrating its performance as an unbiased, discovery-oriented platform for single-cell proteomic analysis of the nervous system. We show that molecularly-guided spatial proteomics enables robust and quantitative profiling of CNS cells across sample fixation conditions, staining modalities, and sampling area, extending to the single-cell regime. A key methodological advantage of this approach is its compatibility with established imaging modalities, including protein-based IF and mRNA-based FISH **(Fig. 1C-D, Supp. Fig. 1B-C)**. Large-scale transcriptomic studies have established mRNA expression as a robust cell-type marker, and in some cases, such as for synaptically localized proteins (e.g., GAD1 in inhibitory neurons), mRNA-based labeling more effectively visualizes neuronal cell bodies than antibody staining, which often preferentially labels distal processes. Using this framework, we successfully resolved brain region-specific neuronal populations **(Fig. 2E)** and identified multiple high-confidence cell-type-specific protein markers **(Fig. 2, Supp. Fig. 4)**, supported by orthogonal histological and transcriptomic ISH evidence.

The increasing availability of well-annotated large-scale single-cell transcriptomic atlases, particularly for mouse and human brain^24,52^, provides a valuable and accessible resource for comparative multi-omics analyses and validation. These atlases can also enable evaluation of analytical strategies in single-cell proteomics. In this study, we leveraged transcriptomic data to assess the impact of different protein data processing approaches and observed that directLFQ-based normalization substantially improved agreement with transcriptomic trends compared to commonly used global imputation and median normalization strategies in these sparse, tissue-derived single-cell conditions **(Fig. 3, A-G).** Moreover, we used transcriptomics to assign confidence in cell-type origin of proteomics hits **(Fig. 3, J, Supp. Fig. 4)**. This refinement is particularly important in heterogeneous tissues such as the brain, where low-input spatial sampling can capture proteins originating from neighboring or intermingled cells and processes. Together, these observations highlight the potential of transcriptomics-informed comparisons as a pragmatic means of benchmarking and refining single-cell proteomics analysis pipelines. Incomplete concordance between transcriptomic and proteomic measurements, consistent with prior studies^34,53,54^, underscores the complementary nature of the two modalities and highlights opportunities for insights from proteomics that are not accessible through transcriptomics alone.

Analysis of neuroimmune responses in the stab wound injury (SWI) model provided additional methodological insights into integrating proteomic and transcriptomic data analyses. We observed substantial overlap between our proteomic hits and a recent SWI transcriptomic study^38^, with comparable fold changes for most shared targets, including Anxa3, B2m, Dhrs1, S100a6, Npl, Ptpn6, Isg15, C1qa, and C1qc. In contrast, CD68, a key inflammatory marker, showed greater enrichment at the protein level (∼6-fold) **(Supp. Fig. S5F)** in our dataset compared to transcriptomic measurements (∼2.5-fold). This disparity likely reflects localized accumulation of CD68 protein near the stab wound injury axis, from which our samples were collected, whereas corresponding mRNA signals are more spatially diffused, as reported previously^38^. Consistent with this, spatial imaging of selected targets in the same study revealed pronounced differences between mRNA and protein localization, further reinforcing that transcriptomic and proteomic measurements provide complementary, rather than interchangeable, information.

Together, these findings highlight the depth and specificity enabled by LCM-based molecularly-guided scSP, positioning this approach as a powerful framework for studying neurodegenerative diseases such as PD, where selective neuronal vulnerability and sparse, mosaic pathology pose persistent challenges to mechanistic understanding. Although SNpc neuronal subtypes have been extensively characterized at the transcriptomic level^42^, their proteomic organization and its relationship to vulnerability remain largely unexplored. Our analysis reveals many proteomic differences between ventral and dorsal SNpc neurons, including alterations in proteins related to dopamine oxidation^55^, such as Aldh1a1 (**Fig. 5, E–F**), suggesting proteomic specializations that may contribute to differential susceptibility, although further work will be required to define the underlying mechanisms.

Within our dataset examining proteomic changes in sparsely populated, aggregate-containing neurons, we observed enrichment of gene ontology terms (**Supp. Fig. 8**) and KEGG pathways (**Supp. Fig. 9**) associated with PD in aggregate-associated differentially expressed proteins (DEPs), supporting the biological robustness of the dataset. These DEPs pointed toward coordinated disruption of proteasomal and mitochondrial pathways (**Fig. 6G**), reflected by widespread alterations in their core subunits, two systems that represent central mechanistic pillars of PD biology^56^. Aggregated αSyn is known to impair proteasomal degradation (McNaught and Jenner 2001, Thibaudeau, Anderson et al. 2018) and directly interacts with mitochondrial membranes^57,58^, leading to respiratory dysfunction. Consistent with this, we observed reduced levels of the PD-associated deubiquitinase Uchl1^59^, indicative of sustained proteostasis failure. Similar proteasomal and mitochondrial alterations were also detected in single-cell proteomics data from an orthogonal PFF-induced PD model in the regenerative spiny mouse^60^, suggesting these responses possibly represent conserved features of αSyn pathology. In parallel, we detected reduced abundance of multiple ribosomal proteins, indicating compromised translational capacity. This aligns with prior reports of ribosomal dysfunction and clinical observations of reduced RPL and RPS subunits in PD patient brains^61,62^, although these studies were conducted at the bulk tissue level. Together, the combined disruption of translation and ubiquitin–proteasome function suggests a potential decoupling between transcript abundance and protein clearance, underscoring the importance of proteomic analyses for accurately defining neuronal state in PD.

Several previous studies have characterized proteomic alterations in PD patient tissue and preclinical models using bulk approaches^63–65^. Bulk and single-cell approaches may exhibit pathway-level convergence; however, the specific proteins underlying these signatures can differ. For example, direct comparison of our DEPs with a recent bulk proteomic study^63^ using a similar experimental paradigm revealed limited overlap, likely reflecting the whole-tissue nature of bulk analysis. In contrast, a more substantial overlap was observed with human Lewy body interactor proteins reported by Killinger et al.^66^, providing validation of our neuron-enriched readouts, given that Lewy bodies are restricted to neurons in PD patients. The limited concordance with bulk datasets is consistent with the inherent non-specificity of whole-tissue measurements, including contributions from non-neuronal cell types and dilution of disease-relevant signals in tissues where pathology is confined to a subset of 1-2% neurons^3^. By resolving proteomes at the level of individual dopaminergic neurons, our single-neuron analysis overcomes limitations from tissue-averaged measurements. Notably, dimensionality-reduction analysis (**Fig. 6D**) revealed a compact cluster of aggregate-free neurons, whereas aggregate-containing neurons exhibited substantial divergence, suggesting heterogeneous stress responses elicited by αSyn pathology that are only resolvable through single-cell approaches and further highlighting the scope of scSP.

Several limitations of the LCM-based scSP approach should be considered, particularly when applied to the brain. First, the complex and intermingled architecture of the brain can lead to proteomic contamination. For example, we detected astrocytic proteins in SNpc neuronal samples **(Fig. 2**, **Fig. 5)**, and microglial proteins in isolated ramified astrocytes in SWI samples **(Fig. 4, Supp. Fig. 5, D-F)**. Contaminating proteins may also originate from closely related cell types, such as different neuronal subpopulations. For instance, in our dopaminergic subtype analysis, Anxa⁻ samples showed significant non-zero proteomic intensity values for Anxa1 **(Fig. 5E)**, unlike the near-zero expression levels of SNpc markers such as TH or Sncg in spatially separated cortical samples **(Fig. 2G)**. This is notable given prior transcriptomic analyses^40^ and IF imaging indicating that Anxa⁻ SNpc neurons exhibit undetectable (not just low) Anxa1 expression **(Fig. 5A, 5F)**. Such effects likely arise from overlapping cells or processes within the target ROI, particularly along the Z-axis, where precise isolation of neuronal cell bodies is inherently challenging. Related to this, proteins predominantly localized to neuronal processes or synapses are underrepresented in single-cell samples compared to larger ROI samples reflecting an intrinsic limitation of cell-body based isolation strategies for cells with complex structures.

Second, sample variability remains a challenge; high coefficient of variations (CVs) and missing values were observed, as is typical for single-cell studies^26,67^. This is particularly important when distinguishing biological or disease-specific differences from variability introduced by methodological heterogeneity. To improve downstream biological interpretability, we purposefully omitted features not present in at least 40% of samples in at least one group (see Methods). While small sample sizes can therefore be sufficient to detect protein differences with large effect sizes (e.g., **Fig. 4**), especially given histology-guided rather than blind isolation of cells, larger sample sizes will likely be required to resolve more subtle differences.

In contexts where cell-specific heterogeneity is driven by biological factors or disease states, disentangling these limitations from technical variability is a challenge, underscoring the need to balance sensitivity and interpretability. Ongoing advances in MS sensitivity, reduced costs for single-cell processing, and continued methodological and analytical improvements in LCM are expected to further enhance the utility of LCM-based scSP for CNS studies. In this study, as discussed above, integration with orthogonal transcriptomic datasets provided additional context to distinguish biologically meaningful heterogeneity from low-input or technical effects. In addition, although not applied here, machine learning-based strategies, such as those used in deep visual proteomics^19,21,49,68^, could further improve cell selection by minimizing overlap among histology-positive neurons and enhancing cell annotation throughput.

In summary, this work establishes single-cell spatial proteomics (scSP) as a powerful approach for studying the nervous system, enabling investigation of diverse biological questions and disease states, and provides a foundation for continued technical and analytical refinement.

## Materials and Methods

### Post-mortem tissue processing

All mouse procedures were approved by the Institutional Animal Care and Use Committee (IACUC) at the California Institute of Technology. Animals were euthanized using a sodium pentobarbital overdose and subjected to transcardial perfusion with chilled PBS. Brains were either collected and flash frozen using a dry ice-ethanol bath, or perfusion was continued with 4% (w/v) paraformaldehyde (PFA) in PBS to fix the brain. Fixed brains were then collected and kept for 24-48 hr in PFA solution and subsequently cryoprotected at 4°C in a solution containing 30% (w/v) sucrose for 72 hr. Brains were flash-frozen in O.C.T. Compound (Scigen, #4586) using a dry ice-ethanol bath and kept at -70 °C until sectioning. Brain sections were obtained using a cryostat (Leica Biosystems) and collected in 1x PBS. Sections were stored at 4°C in PBS (supplemented with 0.02% Azide) for short-term storage or at - 20°C in cryoprotectant (Bioenno Lifesciences, 006799-1L) for longer preservation. Detailed protocol available at 10.17504/protocols.io.14egn1j4yv5d/v1.

### Mouse αSyn PFF preparation

Recombinant mouse αSyn PFFs were produced as previously outlined^60^. In summary, αSyn monomer was produced in E. coli BL21 (DE3) cells utilizing the pT7-7 vector and subsequently purified through size-exclusion chromatography (Cytiva) and anion-exchange chromatography on a HiPrep Q HP 16/10 column (Cytiva). Endotoxin was eliminated using endotoxin-reduction resin (Thermo Fisher Scientific, #88277), resulting in preparations with <0.02 EU/μg. Monomeric αSyn (5 mg/mL in PBS) was filter sterilized with a 0.22 μm filter and incubated at 37 °C with continuous agitation at 1,000 rpm to facilitate fibril formation. Following the elimination of remaining monomer via centrifugation, the resultant fibrils were resuspended in PBS at a concentration of 5 mg/mL, aliquoted into 25 μL portions, and preserved at –80 °C. Prior to injection, fibrils were sonicated with a cup-horn sonicator (Qsonica Q700) to generate short PFF fragments suitable for *in vivo* seeding. Detailed protocol available at 10.17504/protocols.io.14egn1j4yv5d/v1.

### Intracranial stereotaxic surgery

Mice were group housed under controlled conditions, maintained on a 13/11-hour light/dark cycle at ambient temperatures ranging from 71 to 75°F, with humidity levels between 30% and 70%. Stereotaxic surgery was performed as described previously ^60^. Before surgery, animals were anesthetized using isoflurane, and the animal was secured on the stereotaxic frame (Kopf Instruments). For the stab wound injury (SWI) experiment, a needle tip (26 gauge) insertion was performed at the coordinates 0.25 AP, -2 ML, -3 DV and left in place for 5 minutes. For PFF injection, 1.5uL of 5mg/mL sonicated PFF was injected at the coordinates 0.25 AP, -2 ML, -3 DV and PBS was injected into the contralateral hemisphere of the same animal as a control. Following the injection, the scalp was closed using non-absorbable nylon sutures (Ethicon #NW3353). Animals were allowed to recover on a heating pad and monitored for health concerns regularly for the next few days. SWI and PFF-injected animals were culled 7 days and 24 weeks post-injection, respectively, to collect the brain as described above. Detailed protocol available at 10.17504/protocols.io.14egn1j4yv5d/v1.

### Immunofluorescence (IF) staining

For IF staining, 30 µm or 35 µm (**Supp. Fig. 1, B,D**) free-floating brain sections were blocked with 10% (v/v) normal donkey serum in PBS supplemented with 1% Triton-X100 (% v/v) (1% PBST) for 90 min and then incubated with primary antibody solution prepared in 0.3% PBST supplemented with 1% (v/v) normal donkey serum overnight at 4°C. After washing in PBS (3 x 10 min), the sections were incubated with secondary antibodies (in 0.3% PBST with 1% NDS) conjugated with Alexa fluorophores (Jackson ImmunoResearch Laboratories, 1:500 dilution) for 90 min at room temperature and then washed 3 x 10 min in PBS. The tissues were mounted on glass slides or UV-treated metal frame slides with PET membranes (Leica Microsystems, #76463-322) for confocal imaging and laser capture microdissection, respectively. Glass slides were allowed to dry overnight and sealed with a coverslip using mounting media (Thermo Fisher Scientific, #P36970). The following primary antibodies were used for IF staining: anti-tyrosine hydroxylase (EMD Millipore, # AB152, RRID:AB_390204, 1:1000; #MAB318, RRID:AB_2313764, 1:1000); anti-GFAP (Abcam, #ab4674, RRID:AB_304558, 1:1000), anti-Iba1 (FUJIFILM Wako Pure Chemical Corporation, #019-19741, RRID:AB_839504, 1:1000), anti-CD11b (Thermo Fisher Scientific, RRID:AB_467108, #14-0112-82, 1:500), anti-CD68 (Thermo Fisher Scientific, RRID:AB_11151139, #14-0688-82, 1:500), anti-NeuN (Cell Signaling Technology, #94403, RRID:AB_2904530, 1:1000; Abcam, #ab104224, RRID:AB_10711040, 1:1000) , anti-Anxa1(Thermo Fisher Scientific, #71-3400, RRID:AB_2533983, 1:250), anti-Aldh1a1 (Thermo Fisher Scientific, #PA5-17943, RRID:AB_10977372, 1:250), anti-Calb1 (Swanth, #CB38a-200uL, RRID:AB_3107026, 1:500), anti-gamma-synuclein (Genetex, #GTX110483, 1951959, 1:500), anti-Camk2a (Abcam, #ab52476, RRID:AB_868641, 1:500), anti-pSer129-αSyn (EP1536Y) (Abcam, #ab51253, RRID:AB_869973, 1:500), anti-Ubiquitin (Proteintech, #80992-1-RR, RRID:AB_2923694, 1:500), anti-p62/SQSTM1 (Proteintech, #18420-1-AP, RRID:AB_10694431, 1:500). Detailed protocol available at 10.17504/protocols.io.14egn1j4yv5d/v1.

### Hybridization chain reaction (HCR)

Split initiator probes^69^ against *Itpr1, Gad1*, and *Gad2* were designed **(Supp. Table 4)** according to Jang *et al*.^70^ and ordered from Integrated DNA Technologies. All washes and incubations were performed at room temperature and with gentle shaking unless otherwise stated. All wash and incubation buffers were prepared from RNase-free reagents. Free-floating sections of the mouse brain were permeabilized in 1x PBS with 0.1% Triton-X100 for 1 hr. Sections were then incubated for 1 hr at 37 °C in a probe hybridization buffer consisting of 2x SSC, 10 % ethylene carbonate (Sigma-Aldrich, #E26258), and 10% dextran sulfate (Sigma-Aldrich, #3730). Following equilibration in hybridization buffer, the samples were incubated for 16 hr at 37 °C in pre-warmed fresh hybridization buffer plus 2 nM of each probe. After probe hybridization, sections were washed twice for 30 min in stringent wash buffer (2x SSC, 30% ethylene carbonate) at 37 °C, then twice for 30 min in 5x SSC with 0.1 % Tween-20 (Sigma-Aldrich, #P1379), and then incubated in HCR amplification buffer (2x SSC, 10% ethylene carbonate) for 1 hr. Alexa Fluor 647 or 488-conjugated hairpins (Molecular Technologies) were heated to 95 °C for 90 s, then cooled to room temperature for 30 min in the dark. Following equilibration in amplification buffer, the tissue was incubated in amplification buffer with 60 nM hairpins for 16 hr. After HCR amplification, the samples were washed twice for 30 min each in 5x SSC with 0.1 % Tween-20, followed by two washes in 1x PBS for 10 min. Sections were stored in 1x PBS at 4 °C until use. Detailed protocol available at 10.17504/protocols.io.14egn1j4yv5d/v1.

### Confocal Imaging

Images were acquired using (i) a spinning disk confocal microscope (Andor, Oxford Instruments), using a 10x, 0.30 NA air objective or 25x, 0.95 NA water-immersion objective, operated with Fusion software (v2.3.0.44) and visualized using Imaris software(v9.5.1, Oxford Instruments, RRID: SCR_007370) and (ii) a Zeiss LSM 880 (Zeiss), using a 25x, 0.8 NA water-immersion objective or 10x, 0.45 NA air objective, operated with Zen Black (v2.3) and visualized using Zen Blue (v2.5).

### Laser-capture micro-dissection (LCM)

Regions of interest were mounted on PET membranes (Leica Microsystems, #76463-322) and excised using a gravity-driven collection system by a Leica LMD7000 and LMD7. Laser settings are included in **Supp. Table 5**. The cut tissue was collected in a 0.65 mL Low Binding Tube cap (Sorenson SafeSeal, #11300; Eppendorf LoBind, # 0030108434) containing 10 µL of lysis buffer consisting of 50 mM triethylammonium bicarbonate (TEAB) (Thermo Scientific, #90114) and 0.2% n-Dodecyl-beta-Maltoside (DDM) (Thermo Scientific, #89903). Upon collecting, the tube was vortexed upside down for 30 seconds and centrifuged at 13,000 x g for 1 minute at 25°C. The sample was then stored at -80°C until further processing. Detailed protocol available at 10.17504/protocols.io.14egn1j4yv5d/v1.

### Sample preparation for mass spectrometry

Stored samples were thawed and vortexed for 30 seconds in an inverted position with an additional 10 µL of lysis buffer to ensure optimal recovery of the tissue. The sample was then centrifuged at 13,000 x g for 1 minute at 25°C and transferred into a LoBind 384-well PCR plate (Eppendorf, #0030129547). The plate was heated for two hours at 70°C in a QuantStudio^TM^ Real-Time PCR machine with the heated lid set to 85°C for sample lysis and protein denaturation. Following lysis, the plate was centrifuged at 1000 x g for 1 minute and allowed to cool to room temperature. Proteolytic digestion was carried out using a mix of trypsin (Promega, #V5280) and Lys-C (Wako Chemicals, # 125-05061). 1 µL of the digestion mix containing 4 ng of each enzyme was added to each sample. The plate was then incubated at 37°C overnight. Following digestion, samples were centrifuged at 1000 x g for one minute, and digestion was quenched with 0.5 µL of aqueous buffer comprising 2% acetonitrile and 4% formic acid. Detailed protocol available at 10.17504/protocols.io.14egn1j4yv5d/v1.

### LC-MS/MS analysis

Peptides were separated on an Aurora Ultimate UHPLC Column (25cm by 75µm, 1.7µm C18, AUR3-25075C18, IonOpticks) with column temperature maintained at 50°C. 5 µL of each sample was loaded directly onto the analytical column to maximize sensitivity. The LC system (Vanquish Neo UHPLC, Thermo Scientific) was coupled to an Orbitrap Exploris 480 mass spectrometer (Thermo Scientific) with a Nanospray Flex ion source (Thermo Scientific). Peptides were separated using a 60-min gradient at a flow rate of 0.22 µL/min.

For αSyn aggregate-bearing SNpc neurons, the same LC system was coupled to an Orbitrap Astral mass spectrometer (Thermo Scientific) equipped with an EasySpray ion source, with the column maintained at 55°C. These samples were analyzed using a 19.5-min gradient at a flow rate of 0.2 µL/min. Detailed LC gradients for both platforms are provided in the **Supp. Table 6**.

Xcalibur software (Thermo Scientific) was used for method implementation and data acquisition on both mass spectrometers. Data-dependent acquisition (DDA) on the Orbitrap Exploris 480 was performed in positive ion mode (spray voltage 1.6 kV, ion transfer tube at 300°C). Full MS1 scans were acquired over a m/z range of 375-1200 at a resolution of 60,000 with a cycle time of 3 seconds. The maximum injection time was set to auto, and the normalized AGC target was set to 300%. Precursor ions with charges ranging from +2 to +6 were selectively targeted for fragmentation using a minimum intensity threshold of 5e3. Dynamic exclusion was set to exclude after one acquisition, with a 45-second exclusion duration and 10 ppm mass tolerance. MS2 scans were acquired in the Orbitrap at 60,000 resolutions with an isolation window of 1.6 m/z, HCD collision energy set at 28%, and an auto-adjusted maximum injection time. The normalized AGC target was set at 200%.

Data-independent acquisition (DIA) on the Orbitrap Exploris 480 was performed in positive ion mode (spray voltage 1.6 kV, ion transfer tube at 300°C) using parameters adapted from Matzinger *et al.*^23^. Briefly, MS1 full scans were acquired with a range of 375-1200 m/z and a resolution of 120,000. The maximum injection time was set to auto, and the normalized AGC target was set to 300%. MS2 scans were acquired at 60,000 resolutions with a maximum injection time of 118 ms and AGC target of 75%.

For the Orbitrap Astral, DIA was performed in positive ion mode (spray voltage 1.8 kV, ion transfer tube at 275 °C). MS1 spectra were acquired at 240,000 resolution across a scan range of 400-800 m/z with an injection time and an AGC target of 500%. MS2 spectra were acquired on the Astral analyzer with an 80-ms injection time and an AGC target of 800%.

Comprehensive DDA/DIA parameters and mass spectrometer settings for both mass spectrometers are listed in the **Supp. Table 6**. Detailed protocol available at 10.17504/protocols.io.14egn1j4yv5d/v1.

### Mass spectrometry data analysis and statistics

RAW files obtained in DDA mode were processed in Proteome Discoverer 2.5 (Thermo Fisher Scientific) using the SequestHT^71^ search algorithm with Percolator validation. Data was searched against the mouse reference database from Uniprot (SwissProt database, downloaded 27 September 2024, 17232 sequences). Lys-C and Trypsin were selected, allowing for a maximum of two missed cleavages and peptide lengths between seven and 30 amino acids. The mass tolerance for precursor ions was set at 20 ppm, while the fragment mass tolerance was defined as 0.1 Da. INFERYS Rescoring was used in automatic mode, and Percolator was used for search validation based on q-values with a strict false discovery rate (FDR) of 1% at the spectrum level^72,73^.

Carbamidomethylation of cysteine residues was designated as a static modification, and oxidation of methionine residues was considered a dynamic modification. A minimum number of 1 peptide sequence (unique and razor) was required for protein identification. Strict parsimony was used to group proteins. Data was exported from PD2.5 and used for further analysis.

RAW files acquired in DIA mode were processed with DIA-NN (version 2.1) ^74^ using a library-free search with FASTA digest enabled and deep learning–based predictions for spectra and retention times. Default DIA-NN parameters were used unless otherwise specified, with the precursor *m/z* range restricted to 400–800 and a maximum of two missed cleavages allowed.

Precursor FDR cutoff was set to 1.0%. Resulting protein- and peptide-level outputs (in parquet format) were imported into Python for downstream analysis using single-cell analysis packages *scanpy*^75^ and *scpviz*^76^, with additional analysis and visualization performed using general-purpose scientific Python libraries documented in the repository README **(Supp. Table 1)**.

Proteins meeting a stringent confidence threshold (PG.Q.Value < 0.01) were designated as high confidence and retained for subsequent analyses. Additional filtering required proteins to be quantified in at least 40% of samples within at least one group per comparison (e.g., region or condition). Proteins lacking valid gene annotations or exhibiting non-unique abundance profiles were excluded, as isoform-level redundancy combined with sparse peptide support resulted in unreliable quantification. Protein abundances were normalized using directLFQ^30^ which was adopted as the primary normalization strategy. For comparison, an alternative normalization approach involving global imputation to 20% of the minimum observed intensity followed by median normalization was evaluated but not used for downstream analyses. Statistical analysis, sample clustering based on principal component analysis, and volcano plots were generated utilizing scpviz. Kolmogorov-Smirnov (KS) normality test was used to test the distribution for whether proteins unique to certain conditions lay within the original sample distribution (rank plot shown in Fig. 1I). Differences in mean protein and peptide counts between conditions were evaluated using Student’s *t*-test. Protein abundance was plotted using GraphPad Prism (v9.5.1, http://www.graphpad.com, RRID: SCR_002798).

### Transcriptomics data analysis and multi-omics comparison

Single-cell RNA sequencing (scRNA-seq) data were obtained from the Allen Brain Cell Atlas mouse whole-brain cell-type atlas (10x Genomics v3) ^24,31^. Data processing and quality control were performed as previously described by Yao et al.^31^, including stringent filtering based on gene detection thresholds, quality control metrics, and doublet scores. The resulting curated scRNA-seq datasets were used directly for downstream analyses. Processed datasets from WMB-10Xv3-Isocortex-1, WMB-10Xv3-Isocortex-2, and WMB-10Xv3-MB were combined, with cells selected based on supertype annotations corresponding to cortical (“CTX”) and midbrain dopaminergic populations (“*-VTA”). Additional filtering retained cells whose region-of-interest acronyms matched MOp (motor cortex) or MB (midbrain). Cells annotated as “L5 NP CTX Glut” and “SNr” were excluded, as these populations do not correspond to the neuronal subtypes analyzed in the proteomics experiments. Regions were annotated by mapping MOp to cortex and MB to SNpc. Transcriptomic data were converted to AnnData format and analyzed using Scanpy, including dimensionality reduction and visualization using uniform manifold approximation and projection (UMAP).

For multi-omics analysis, proteomics data were imported and processed using scpviz as described above. The protein-level AnnData object (.prot) from the pAnnData structure was used to enable direct integration with transcriptomic analyses. Differentially expressed proteins (DEPs) and genes were identified using a fold-change threshold of at least two and a significance cutoff of p < 0.05 based on Student’s t-test. Concordance between transcriptomic and proteomic changes was assessed by comparing the directionality of differential expression, categorizing features as consistently upregulated, consistently downregulated, or oppositely regulated across the two modalities.

To assess the cell-type origin of DEPs, we queried corresponding genes against Drop-seq transcriptomic datasets using the DropViz portal. DEP gene lists were entered as query genes, and the relevant brain region was selected as either the frontal cortex or substantia nigra, as appropriate. Major cell classes were preselected, including astrocytes, neurons, oligodendrocytes, endothelial cells, and microglia/macrophages. When multiple clusters were present within a given cell class, the cluster most relevant to the experimental context was selected (e.g., SN Neuron_Th [#4] for SNpc dopaminergic neurons and FC Neuron_Layer5_Parm1 [#7] for cortical neurons). Natural log–transformed expression values were extracted and plotted in correlation plots using GraphPad Prism.

### Bioinformatics and enrichment analyses

Gene ontology (GO) analysis comparing fixed and frozen tissue was performed using string-db^77^. For the αSyn PFF experiment, pathway enrichment analysis was conducted using ShinyGO (v0.85.1; http://bioinformatics.sdstate.edu/go/, RRID:SCR_019213) with Kyoto Encyclopedia of Genes and Genomes (KEGG; http://www.kegg.jp, RRID:SCR_012773) as the reference database.

Analyses were performed for mouse genes (GRCm39), applying an FDR cutoff of 0.05, pathway size limits of 5–5,000 genes, and reporting the top 10 enriched pathways. Enrichment results were visualized as hierarchical enrichment trees and KEGG pathway maps.

GO enrichment analysis was additionally performed using Metascape (https://metascape.org, RRID:SCR_016620), with annotations spanning biological process (BP), molecular function (MF), and cellular component (CC). Enrichment was assessed using a minimum overlap of three genes, a P-value cutoff of 0.01, and all genes as background. GO term visualization was carried out using Cytoscape (v3.10.3; https://cytoscape.org, RRID:SCR_003032).

Protein–protein interaction (PPI) networks were generated using STRING (https://string-db.org, RRID:SCR_005223), using the full STRING network with experimentally supported interactions only, highest confidence threshold (0.9), and MCL clustering.

## Supporting information

Supp. Table 1_KeyResources_v2

Supp. Table 2_OmicsComparison

Supp. Table 3_dropviz data_SNpc_Cortex

Supp. Table 4_RNA-FISH-probes

Supp. Table 5_LCMSettings

Supp. Table 6_LCMSMSSettings

## Acknowledgments

This research was funded in part by Aligning Science Across Parkinson’s (ASAP-020495 to V.G.) through the Michael J. Fox Foundation for Parkinson’s Research (MJFF), the NIH R01MH136394 to M.L.R and T-F. C., Wellcome Leap Fellowship (through its Delta Tissue program) to M.L.R., and A*STAR National Science Scholarship (BS-PhD) to M.P. V.G. is an Investigator of the Howard Hughes Medical Institute. For the purpose of open access, the authors have applied a CC BY public copyright license to all Author Accepted Manuscripts arising from this submission.

We thank the CNSI Advanced Light Microscopy/Spectroscopy Facility at UCLA and Leica Biosystems for access to the LCM instruments.

## Author contributions

S.D. conceived the overall study. S.D. and M.P. designed the experiments, performed LCM sample collection, data analysis, and visualization, and prepared the manuscript with input from all authors. S.D. conducted stereotaxic surgery, tissue processing, immunohistology staining, Drop-seq analysis, and imaging. M.P conducted proteomics sample preparation, mass spectrometry data acquisition, and all proteomics data processing and analysis, as well as single-cell transcriptomics data analysis. G.M.C. designed and processed samples for RNA-FISH experiments. S.G. assisted with tissue processing, staining, and data visualization plots. M.L.R. advised on single-cell proteomics experiments and instrumentation. T-F.C. supervised proteomics experiments and data analysis. V.G. supervised the overall study and advised on experimental design.

## Data Availability

The data, code, protocols, and key lab materials used and generated in this study are listed in a Key Resource Table alongside their persistent identifiers (**Supp. Table 1**). The proteomics datasets are available on MassIVE MSV000100704 or ProteomeXchange PXD074008.

## Code Availability

All analysis scripts used in this study at available at https://github.com/gnaprs/scSP_CNS-preprint.

