## supplementary figures for "Molecularly-guided spatial proteomics captures single-cell identity of the healthy and diseased nervous system"

### Supplementary figure 1

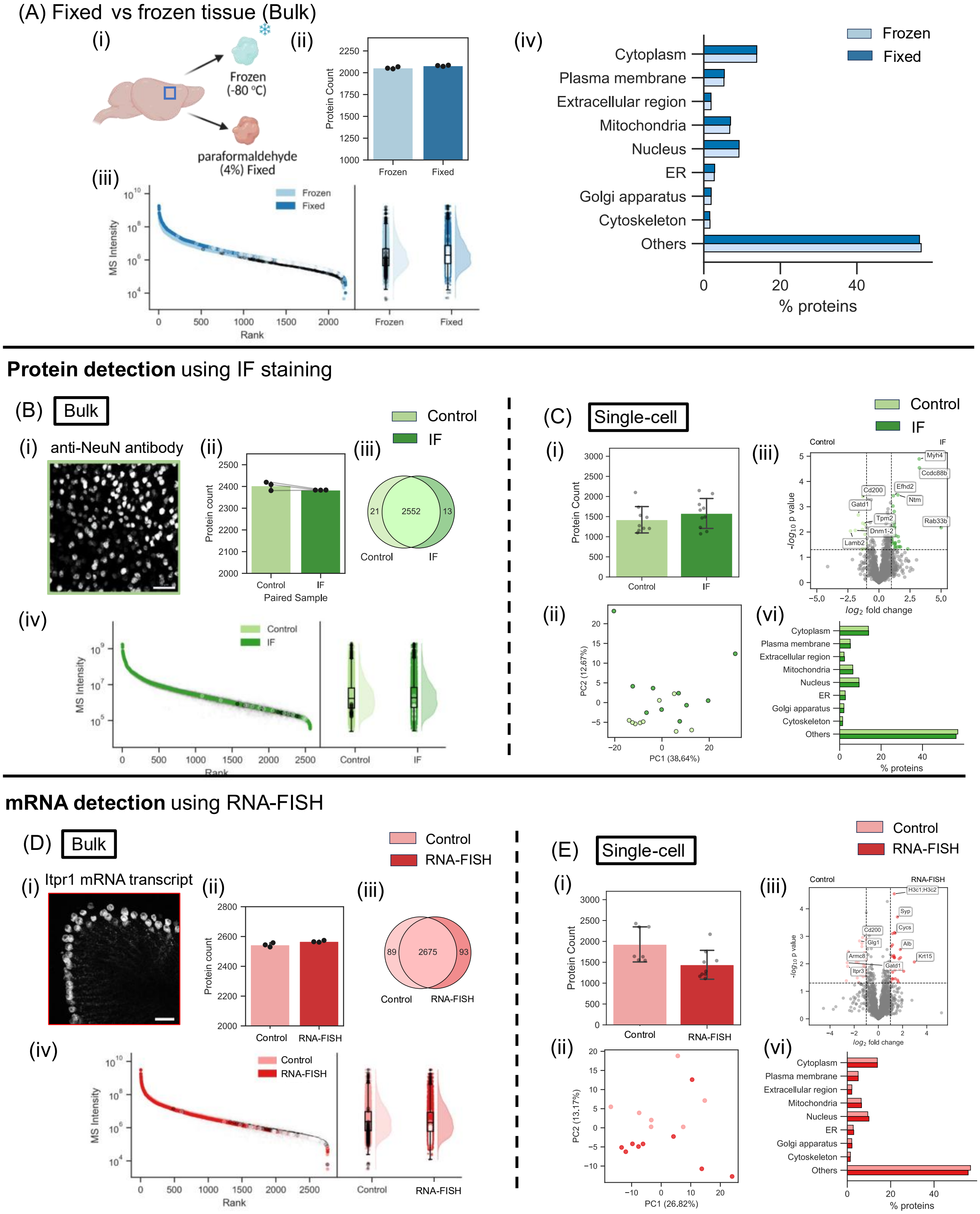

**Supplementary figure 1: Evaluation of fixation and staining effects on proteomic coverage in bulk CNS tissue. (A) Comparison of proteomes from fresh-frozen and 4% PFA-fixed cortical tissue.** (i) Schematic illustrating isolated frozen (-80°C) and 4% PFA-fixed CNS tissue used in this analysis. Created in BioRender. Dutta, S. (2026) https://BioRender.com/otcc3mo.(ii) Protein counts for each condition. (iii) Rank–quantile plot with raincloud analysis highlighting proteins uniquely identified in frozen (black points) or fixed (white points) samples. (iv) Distribution of cellular components derived from selected GO cellular component annotations for proteins detected in frozen and fixed tissue. Values indicate the fraction of proteins assigned to each compartment relative to the total number of quantified proteins. **(B) Effects of immunofluorescence (IF) staining on protein for bulk samples:** (i) Representative anti-NeuN IF image. Scale bar: 50 µm (ii) Protein counts for IF-stained samples and paired PBS controls from the unstained hemisphere of the same mouse (n = 3 animals). (iii) Overlap of protein identifications between the groups. (iv) Rank–quantile plot with raincloud analysis showing uniquely identified proteins in control (black points) and IF-stained (white points) samples. **(C) Effects of IF staining on protein for single-cell samples:** (i) Protein counts for stained and control cells. (n= 10 cells for IF-stained; 9 cells for control) (ii) UMAP projection of protein abundance profiles showing clustering by staining condition. (iii) Volcano plot of differentially abundant proteins between IF-stained and control cells. (iv) Distribution of cellular components derived from selected GO cellular component annotations for proteins detected in IF and control tissue. Values indicate the fraction of proteins assigned to each compartment relative to the total number of quantified proteins. **(D) Fluorescence In-situ hybridization (FISH) for mRNA staining (RNA-FISH) for bulk samples:** (i) Representative RNA-FISH image of cerebellum showing hybridization against the *Iptr* transcript. Scale bar: 50 µm. (ii) Protein counts for RNA-FISH-stained samples and unstained PBS controls (n = 3 animals). (iii) Overlap of protein identifications between the groups. (iv) Rank–quantile plot with raincloud analysis showing uniquely identified proteins in control (black points) and RNA-FISH-stained (white points) samples. **(E) RNA-FISH for single-cell samples:** (i) Protein counts for RNA-FISH-stained and control cells. (n= 7 cells for RNA-FISH-stained; 9 cells for control) (ii) UMAP projection of protein abundance profiles showing clustering by staining condition. (iii) Volcano plot of differentially abundant proteins between RNA-FISH-stained and control cells. (iv) Distribution of cellular components derived from selected GO cellular component annotations for proteins detected in RNA-FISH and control tissue. Values indicate the fraction of proteins assigned to each compartment relative to the total number of quantified proteins

**Supplementary figure 2**

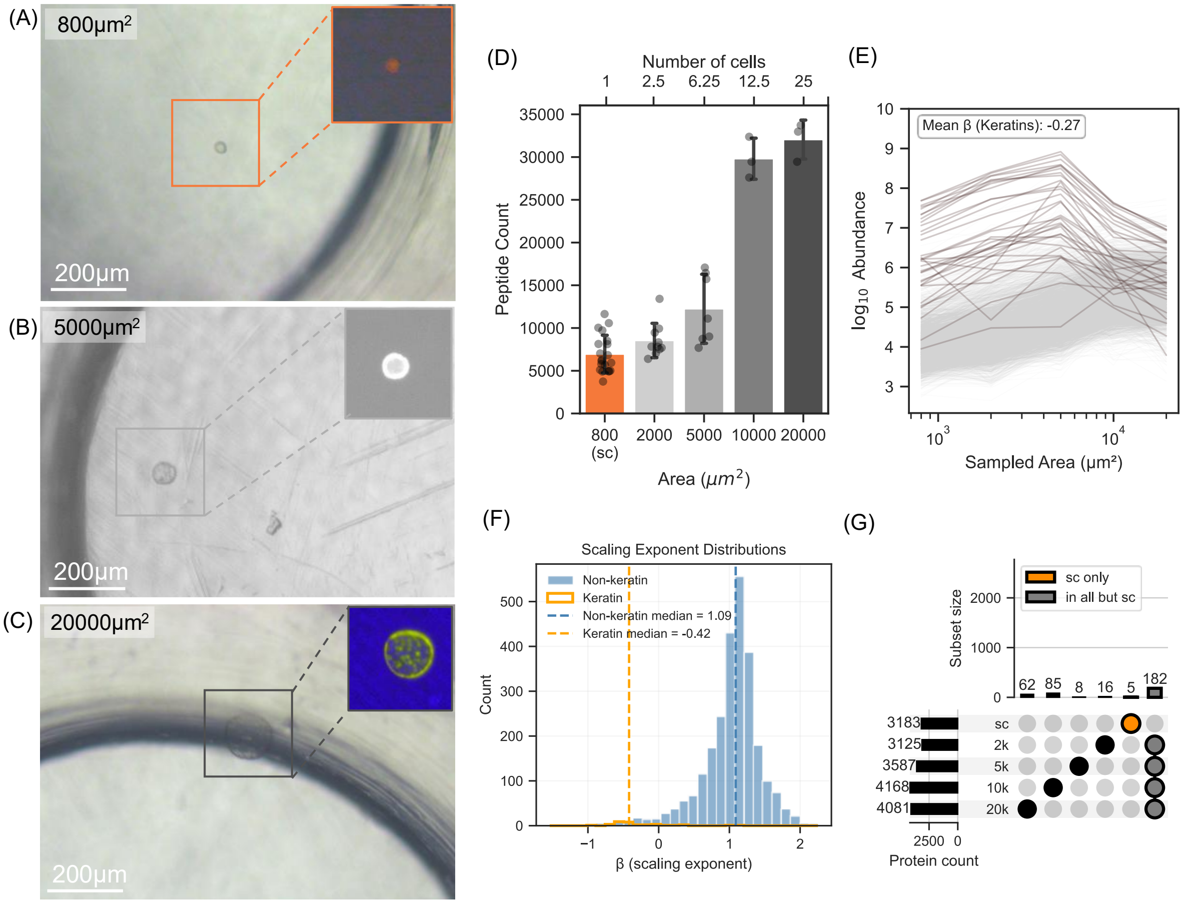


**Supplementary figure 2. Visual confirmation of brain micro-tissue samples after ROI collection and evaluation of input-size scaling in LCM spatial proteomics samples.** **(A)** Representative image of a 20,000 μm² tissue section collected in a 0.5mL tube cap (inset: fluorescence image, 488 channel) **(B)** Representative image of a 5000 μm² tissue section collected in a tube cap (inset: fluorescence image, Cy3 channel) **(C)** Representative image of an 800 μm² tissue section collected in a tube cap (inset: fluorescence image, Cy5 channel) **(D)** Number of high-confidence peptides (q<0.01) identified across sampled areas. **(E)** Log–log scaling of protein abundance as a function of sampled area, with keratin contaminants highlighted in brown. Keratin proteins exhibit a negative scaling exponent ($\beta_1 \approx -0.27$), indicative of variability arising from sample preparation artifacts rather than biological effects. **(F)** Distribution of scaling exponents for non-keratin and keratin proteins. Non-keratin proteins display a median $\beta_1$ of 1.09, consistent with linear scaling with sampled area, whereas keratin proteins show a median $\beta_1$ of –0.42, forming a distinct distribution. **(G)** UpSet plot illustrating protein intersections across sampled areas. 5 proteins were uniquely detected in single-cell samples, and 182 proteins were identified in all area sizes except single-cell samples.

**Supplementary figure 3**

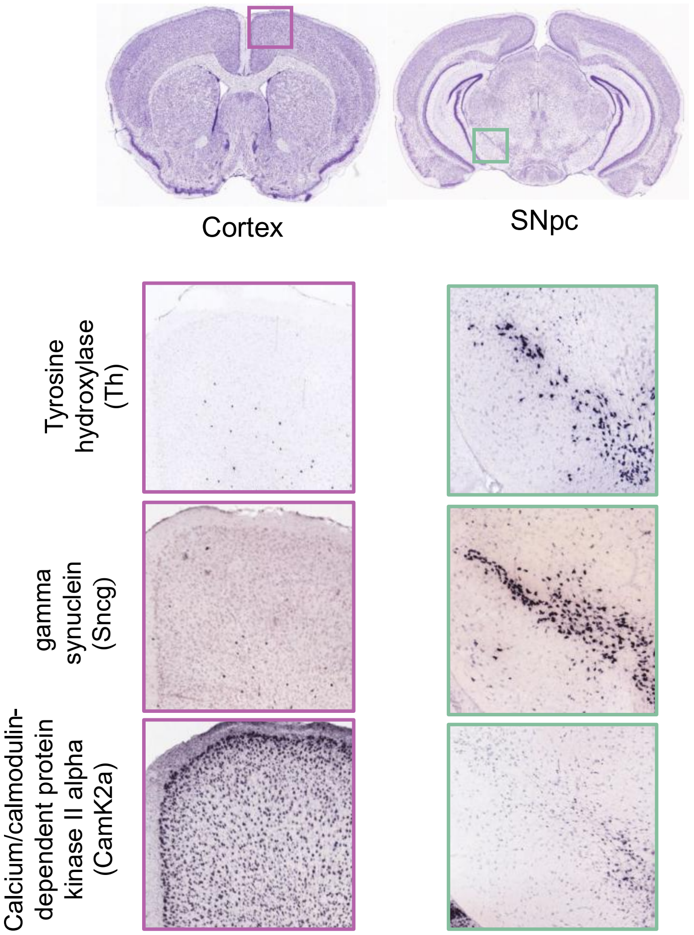


**Supplementary figure 3: mRNA expression levels from the Allen Brain Atlas (ABA) of representative marker proteins identified in the SNpc and cortex:** The magenta and green boxes on the Nissl-stained reference coronal brain section image indicate the approximate locations of the zoomed-in cortical and SNpc panels, respectively. Spatial *in situ* hybridization (ISH) data from the ABA illustrates the mRNA distribution for tyrosine hydroxylase (TH), gamma synuclein (Sncg), and calcium/calmodulin-dependent protein kinase II alpha (Camk2a) (sourced from the mouse coronal ISH atlas published by the Allen Brain Initiative; TH-https://mouse.brain-map.org/experiment/show/1056; Sncg- https://mouse.brain-map.org/experiment/show/72081426; Camk2a-https://mouse.brain-map.org/experiment/show/79490122).

**Supplementary Figure 4**

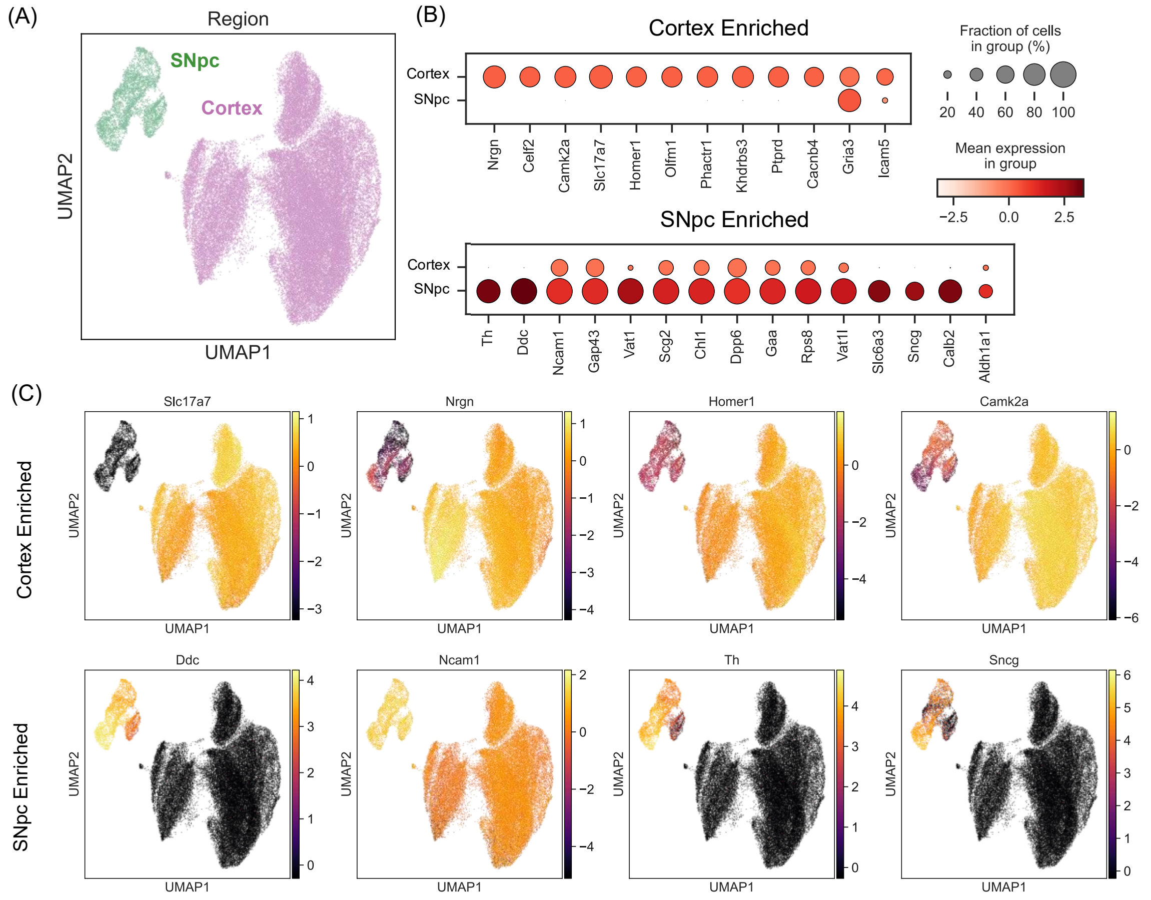


**Supplementary Figure 4: Transcriptomic validation of region-specific neuronal markers using Allen Brain Atlas single-cell RNA-seq data. (A)** UMAP embedding of neurons from the Allen Brain Institute mouse whole-brain 10x scRNA-seq dataset, showing clear segregation of cortical neurons (IT–ET glutamatergic neurons from MOp; purple) and midbrain dopaminergic neurons corresponding to the substantia nigra pars compacta (SNpc; green). (n= 4154 cells for SNpC; 48719 cells for cortex) **(B)** Dot plot summarizing the top ten and some selected genes enriched in cortical neurons (top) and SNpc neurons (bottom) with dot size indicating the fraction of cells expressing each gene and color intensity representing mean scaled expression. **(C)** Feature-level UMAP projections illustrating selective examples of cortical-enriched marker expression, including Slc17a7, Nrgn, Homer1, and Camk2a, and SNpc-enriched marker expression, including Ddc, Ncam1, Th, and Sncg.

**Supplementary figure 5:**

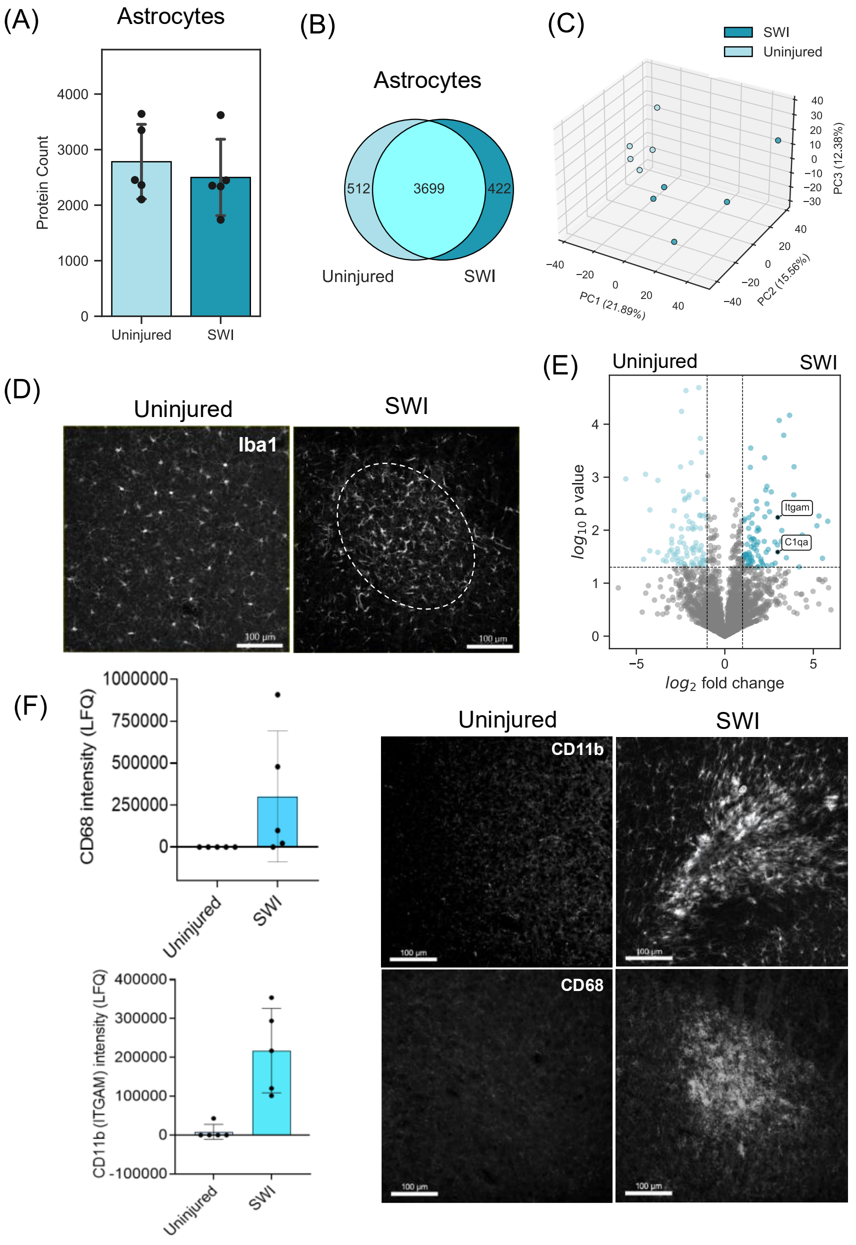


**Supplementary figure 5. Spatial proteomics in stab wound injury (SWI) samples identifies reactive microglial markers at the site of acute injury: (A)** Protein group counts per GFAP$^+$ astrocyte isolated from the contralateral uninjured hemisphere and the peri-lesional region following SWI. **(B)** Shared and condition-specific protein groups detected in astrocytes from uninjured and SWI tissue. **(C)** Three-dimensional PCA of astrocytic proteomes demonstrating separation between uninjured and SWI samples. **(D)** Representative IF images of Iba1-positive microglia in the uninjured site and at the SWI site, showing injury-induced microglial activation and morphological remodeling. Scale bars, 100 μm. **(E)** Volcano plot depicting representative microglial proteins detected in the SWI samples. **(F)** LFQ intensities of the example microglial activation markers integrin alpha M (ITGAM/CD11b) and CD68 in control and SWI samples, with corresponding IF validation. Scale bars, 100 μm.

**Supplementary figure 6:**

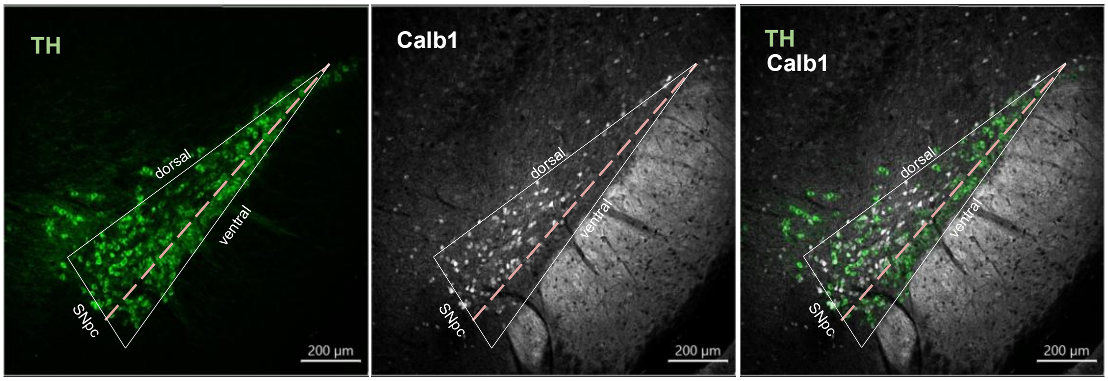


**Supplementary figure 6: Dorsal SNpc neurons are positive for Calb1.** IF characterization of TH neurons confirms Calb1+ dopaminergic neurons are specifically located in the dorsal SNpc.

**Supplementary figure 7:**

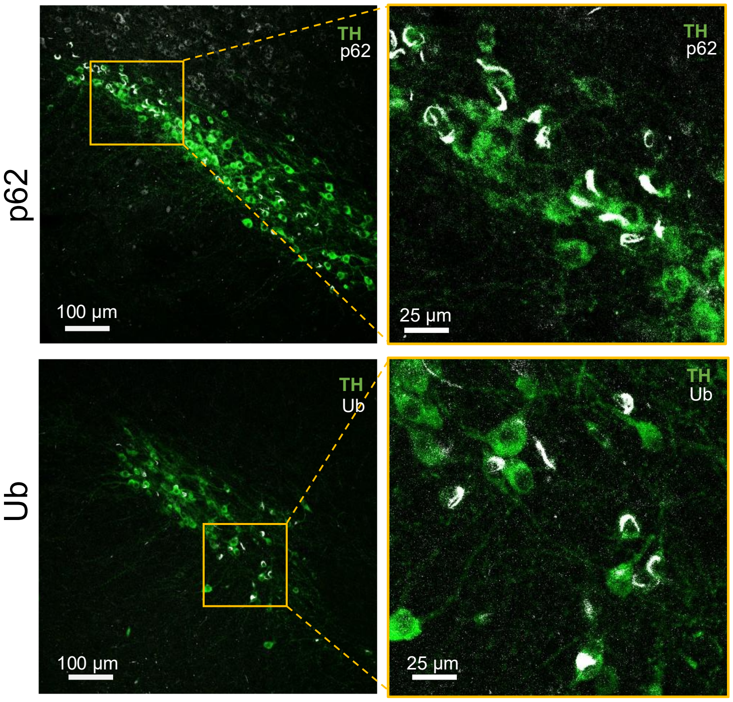


**Supplementary figure 7: PFF-induced intraneuronal aggregates are positive for orthogonal pathological markers of PD.** IF characterization of SNpc neurons following PFF injection shows co-labeling of canonical intra-neuronal aggregation markers ubiquitin and p62.

**Supplementary figure 8**

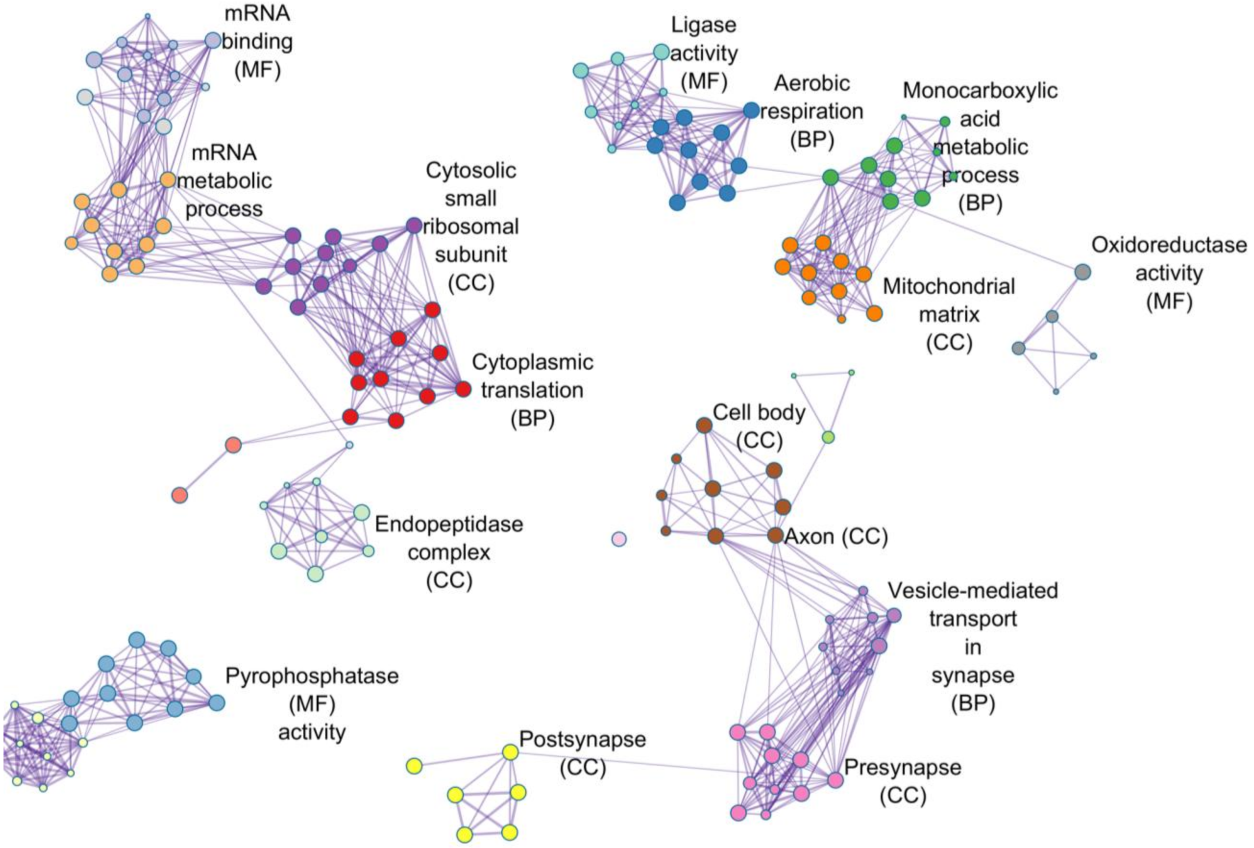


**Supplementary figure 8. Gene ontology enrichment of differentially abundant proteins in aggregate-bearing neurons.** Gene ontology analysis of proteins altered in $Agg^{+}$ versus $Agg^{-}$ neurons reveals enrichment across cellular component (CC), molecular function (MF), and biological process (BP) categories consistent with neurodegenerative pathology.

### Supplementary figure 9

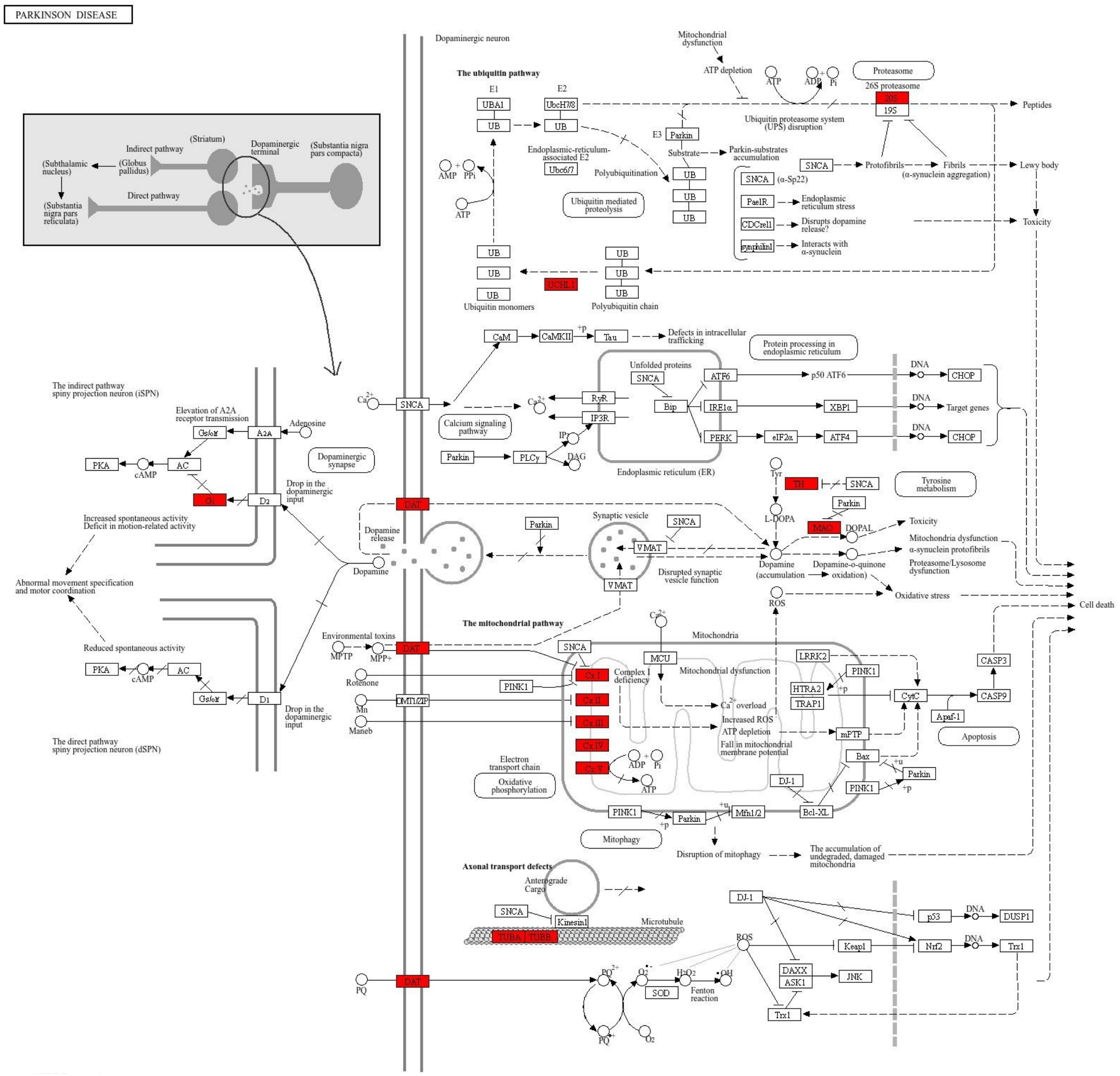


**Supplementary Figure 9. Differentially regulated proteins in Agg+ neurons mapped onto the Parkinson's disease pathway.** Proteins that are up- or down-regulated in Agg+ neurons and are key components of the Parkinson's disease pathway, as defined by the Kyoto Encyclopedia of Genes and Genomes (KEGG), are highlighted in red. These mapped proteins span multiple pathogenic nodes, including dopaminergic signaling, mitochondrial dysfunction, the ubiquitin-proteasome system, calcium signaling, axonal transport, and oxidative stress, illustrating widespread perturbations within established molecular pathways relevant to Parkinson's disease.

**Supplementary figure 10.**

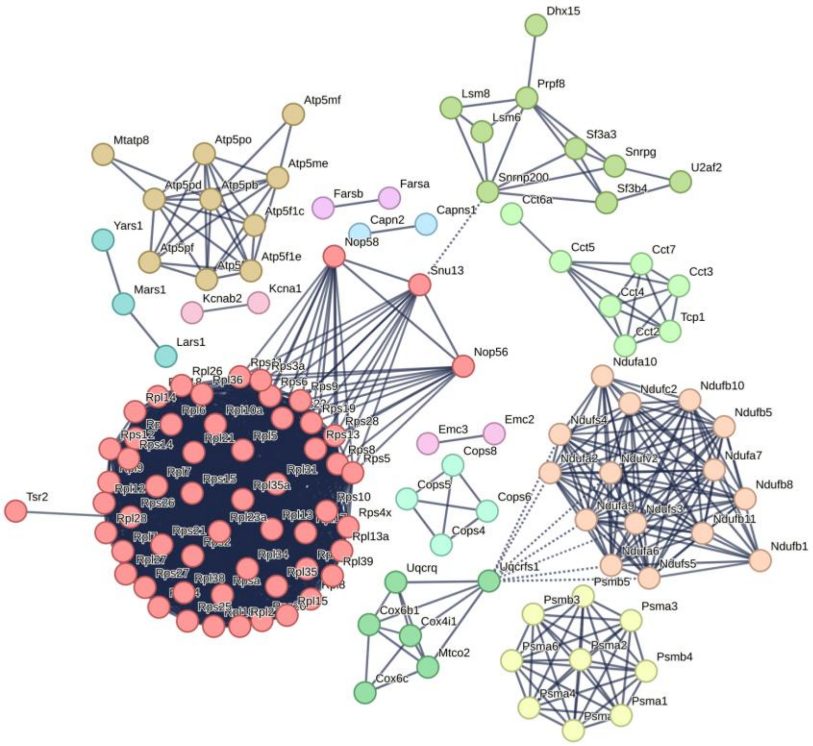


| color | cluster Id | gene count | description |
|---|---|---|---|
| red | Cluster 1 | 57 | + Cytoplasmic translation |
| peach | Cluster 2 | 15 | + Proton motive force-driven mitochondrial ATP synthesis |
| tan | Cluster 3 | 10 | + Formation of ATP by chemiosmotic coupling |
| yellow | Cluster 4 | 9 | + Proteasome |
| light green | Cluster 5 | 9 | + Spliceosome |
| pale green | Cluster 6 | 7 | + Positive regulation of establishment of protein localization to telomere |
| green | Cluster 7 | 6 | + Cardiac muscle contraction |
| mint | Cluster 8 | 4 | + Protein deneddylation |
| teal | Cluster 9 | 3 | + Aminoacyl-tRNA biosynthesis |
| light blue | Cluster 10 | 2 | Calpain complex |
| lilac | Cluster 11 | 2 | phenylalanine-tRNA ligase complex |
| pink-violet | Cluster 12 | 2 | + Protein insertion into ER membrane by stop-transfer membrane-anchor sequence |
| pink | Cluster 13 | 2 | + Juxtaparanode region of axon |

**Supplementary Figure 10. Protein–protein interaction network of DEPs in Agg$^{+}$ neurons.** Nodes represent individual proteins and edges represent known or predicted interactions. Node colors indicate functional clusters, including cytoplasmic translation (Cluster 1, red), mitochondrial ATP synthesis (Clusters 2–3, peach/light orange), proteasome (Cluster 4, yellow), spliceosome (Cluster 5, light green), and other modules (Clusters 6–13, varying colors as shown). The network highlights tightly connected modules of ribosomal and RNA-binding proteins, including spliceosome components, reflecting coordinated dysregulation in aggregate-bearing neurons.
